# Multivariate classification of multichannel long-term electrophysiology data identifies different sleep stages in fruit flies

**DOI:** 10.1101/2023.06.12.544704

**Authors:** Sridhar R. Jagannathan, Rhiannon Jeans, Matthew N. Van De Poll, Bruno van Swinderen

## Abstract

Sleep is observed in most animals, which suggests it subserves a fundamental process associated with adaptive biological functions. However, the evidence to directly associate sleep with a specific function is lacking, in part because sleep is not a single process in many animals. In humans and other mammals, different sleep stages have traditionally been identified using electroencephalograms (EEGs), but such an approach is not feasible in different animals such as insects. Here, we perform long-term multichannel local field potential (LFP) recordings in the brains of behaving flies undergoing spontaneous sleep bouts. We developed protocols to allow for consistent spatial recordings of LFPs across multiple flies, allowing us to compare the LFP activity across awake and sleep periods and further compare the same to induced sleep. Using machine learning, we uncover the existence of distinct temporal stages of sleep and explore the associated spatial and spectral features across the fly brain. Further, we analyze the electrophysiological correlates of micro-behaviours associated with certain sleep stages. We confirm the existence of a distinct sleep stage associated with rhythmic proboscis extensions and show that spectral features of this sleep-related behavior differ significantly from those associated with the same behavior during wakefulness, indicating a dissociation between behavior and the brain states wherein these behaviors reside.

## Introduction

Humans spend a third of their life engaged in sleep, wherein they become less responsive to external stimuli. Most animals studied so far, starting from the tiny fruit fly to the large sperm whale (Miller et al. 2008) display extended periods of quiescence, which are now categorized as sleep. Evolutionary conservation of the sleep state in all animals suggests that its benefits outweigh the potential risks and vulnerabilities brought on by losing awareness of one’s external environment. Sleep deprivation has been shown to produce deficits in learning and memory (Rasch and Born 2013), immune system malfunction (Besedovsky, Lange, and Born 2012) and stress regulation (Paul J. Shaw et al. 2002). However, the organization of sleep in relation to its potential functions remains unclear. Different theories have been proposed for functions of sleep including those involving processes like neuronal plasticity and synaptic downscaling (Cirelli and Tononi 2008) and metabolic waste clearance (Xie et al. 2013). However, sleep research methodology is largely driven by research in humans and other mammals and the primary way of classifying sleep states has therefore been using electrophysiological readouts, such as electroencephalography (EEG). By identifying distinct electrical signatures associated with the different stages of sleep, different functional roles have been hypothesized for them. For example, rapid eye movement (REM) sleep in mammals has been proposed to regulate motor learning and memory consolidation (Siegel 2001; Walker and Stickgold 2004), while slow wave sleep (SWS) has been proposed to regulate synaptic strength and homeostasis mechanisms (Tononi and Cirelli 2014).

One of the primary challenges for understanding sleep architecture has been developing a capacity to record and assess brain-wide patterns of electrical activity across long time periods that encompass several sleep-wake transitions. In this context, small animals such as the fruitfly *Drosophila melanogaster* present as extremely challenging subjects, even though they could potentially provide a wealth of molecular genetic tools to help better understand sleep biology. Previous sleep studies in flies have either recorded from just a single LFP channel during spontaneous sleep bouts (Yap et al. 2017; van Alphen et al. 2013; Nitz et al. 2002; B. van Swinderen, Nitz, and Greenspan 2004), or from multi channel probes during short (∼15min) bouts of genetically-induced sleep (Yap et al. 2017; Paulk et al. 2013). In other work, whole-brain calcium imaging in sleep-deprived flies revealed distinct stages of spontaneous sleep (Tainton-Heap et al. 2021), although these recordings were rarely long enough to display any revealing sleep architecture, and it remains unclear how these different sleep stages might be manifested across the fly brain from the central complex to optic lobes.

The primary reasons for the lack of whole-brain or multichannel sleep data in *Drosophila* are technical in nature: a) it is difficult to perform long-term electrophysiological recordings with multiple electrodes in such small brains; the survival rate is low; and the recording tools used do not yet allow for consistent spatial positioning of multiple electrodes across different flies. b) calcium imaging on the other hand, which lacks in temporal precision compared to LFPs, does allow for consistent spatial locations of recordings (with image registration tools), however concerns with photobleaching and phototoxicity have made it difficult to achieve the long-term recordings to acquire spontaneous sleep data. Subsampling provides one solution: for example, in a recent study 24 hr recordings were conducted by recording for only 1 sec after every minute (thus recording for only 1.6% of the overall time) (Flores-Valle and Seelig 2022). However, this subsampling approach might miss important sleep transitions or longer-lasting sleep phenomena. To best compare the brain activity during sleep in flies with similar data from other animals would ideally involve similar readouts akin to a whole-brain EEG, which in *Drosophila* would necessarily involve miniaturized multichannel probes such as used previously for visual studies (Paulk et al. 2013, 2015) as well induced sleep (Yap et al. 2017) and anesthesia experiments (Leung et al. 2021; Cohen, van Swinderen, and Tsuchiya 2018; Cohen et al. 2016; Cohen and Tsuchiya 2018; Muñoz et al. 2020). Additionally, such recordings would ideally be supplemented by detailed behavioral analysis beyond the simple locomotory determinants that have traditionally defined sleep in flies (P. J. Shaw et al. 2000; Hendricks et al. 2000). Mammalian sleep stages involve distinct micro-behaviors in addition to electrophysiological correlates (Dement and Kleitman 1957; Fulda et al. 2011), and this seems to be true for invertebrates as well (van Alphen et al. 2021; Rößler et al. 2022; Iglesias et al. 2019).

In this study, we optimized a multichannel LFP recording preparation for *Drosophila* flies, to track long-term neural activity in 16 channels across one hemisphere of the fly brain, in a transect from the retina to the central complex. The flies underwent spontaneous sleep bouts while walking/resting on an air-supported ball, and survived long enough to provide 20 hrs of data over one circadian cycle. We developed calibration tools to consistently record from similar spatial locations in different flies. We used machine learning based methods (support vector machines and random forest classifiers) to first investigate the structure of sleep bouts, and further explored the spectral features across multiple brain channels. We also employed machine learning techniques (pose tracking and identification) to identify fly micro-behaviors during these long-term recordings, to determine their potential association with different sleep stages. Taken together our analyses identify distinct sleep stages in the fly central brain, with rhythmic proboscis extensions being a key behavioral feature. We find that the LFP features associated with proboscis extensions during wake and sleep are dissimilar, suggesting that a distinct brain state is driving the sleep functions associated with this rhythmic micro-behavior.

## Results

### Behavioral analysis of tethered flies during sleep and wake

Prior to conducting any electrophysiological recordings, we first investigated how flies slept when tethered to a rigid metal post while being able to walk on an air-supported ball (*Figure 1A*). Flies were filmed overnight under infrared illumination, and locomotory behavior was quantified using a pixel subtraction method (Yap et al. 2017) to identify sleep epochs, defined by the absence of locomotion or grooming behavior for 5 minutes or more (P. J. Shaw et al. 2000; Hendricks et al. 2000; van Alphen et al. 2013; Yap et al. 2017). We also tracked the movement of different body parts, including the proboscis, antennae, and abdomen to detect potential micro-behaviors during sleep. For this, we used machine learning (DeepLabCut (Mathis et al. 2018)) to train a classifier to track micro-behavioral movements through wake as well as sleep (*Figure 1B*). As shown previously (Yap et al. 2017) tethered flies were able to sleep in this context (*Figure 1C; Figure S1A*). As described recently (van Alphen et al. 2021), we also observed regular proboscis extensions (PEs) during wake, as well as during sleep bouts (*Figure S1B*), which often occurred in rhythmic succession (*Figure 1C, orange trace*). We also observed antennal movements and were surprised to discover that these were oscillatory in a subset of flies (*Figure 1C, red trace*). Since PEs were also often rhythmic during sleep, we characterized both micro-behaviors in the frequency domain (*Figure 1D,E, top*) to determine if these were different between sleep and wake. We found that a greater proportion of the sleeping states displayed both antennal periodicity as well as PE periodicity, compared to the waking states (*Figure 1D,E, bottom*), and that antennal periodicity occurred at a small but significantly lower frequency during wake (*Figure S1G*). However, the time course and presence of individual proboscis extensions (*Figure S1B/C*), as well as the dynamics (e.g., periodicity, frequency) of periodic proboscis extensions were not different between sleep and wake (*Figure S1F*), even if this presence varied across sleep and wake.

**Figure 1:**
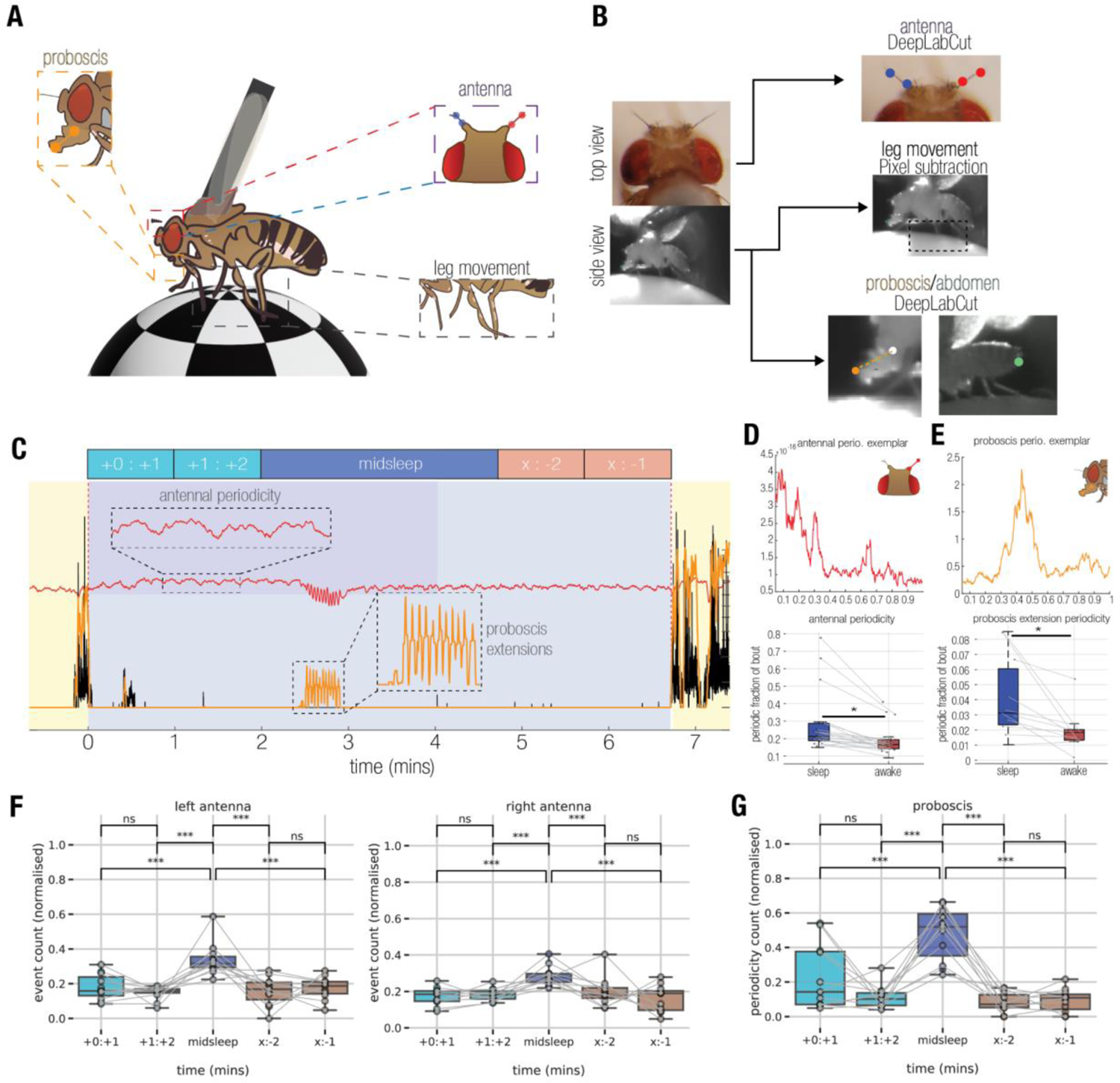
Micro-behaviors of tethered flies. **A)** Schema for the setup used to record micro-behaviors of sleeping and waking flies. A tethered fly stands on an air-supported ball. **B)** The fly is filmed by two cameras. Footage from these cameras is fed through a preprocessing pipeline that tracks movements of the antennae (Top), legs (Middle), abdomen and proboscis (Bottom). **C)** An example sleep bout from a fly. Locomotive activity (Black) has ceased for long enough that the period of inactivity is classified as a sleep bout. The movement of the right antenna (Red trace) shows an apparent low frequency periodicity (See inset) across the 6 minute bout, interrupted in the middle by a series of proboscis extensions (Orange trace; See inset). **D)** Top, FFT of antennal activity during an exemplar sleep bout containing antennal periodicity. Bottom, Comparison of the fraction of sleep and wake that consisted of periodic antennal activity (*p < 0.05; Student’s T-test). **E)** As with D, for proboscis periodicity. **F)** Proportions of antennal periodicity (left and right antennae) across different sleep segments: +0:+1 indicates 1 mins after start of sleep, +1:+2 indicates 2 mins after start of sleep, x:-2 indicates 2 mins before end of sleep, x:-1 indicates 1 mins before end of sleep. The normalized count is significantly higher in the midsleep segments compared to other segments. **G)** As with F, but for periodic extensions of the proboscis***p<0.001, ns indicates not significant.

A previous study suggested that PEs during sleep are accomplishing a specific function in flies linked to waste clearance, and that these might be specific to a deeper sleep stage (van Alphen et al. 2021). We therefore next examined if PE and antennal periodicity varied throughout a sleep bout. For this, we segmented all >5 min sleep bouts into 5 distinct epochs, as done previously for spontaneous sleep experiments in tethered flies (Yap et al. 2017; Tainton-Heap et al. 2021) (*Figure 1C, top schema*). The first 2 and last 2 minutes of sleep (flanked by locomotory behavior) were analyzed separately for micro-behaviors, and compared to ‘midsleep’ epochs which could be of different durations. To understand if likelihood of periodicity for both antennae and proboscis vary based on the sleep epochs we used multilevel modeling instead of traditional repeated measures of analysis of variance (as different flies had varying numbers of sleep epochs). For details refer to the methods section (Multilevel models - models for antennal, proboscis periodicity). For all the micro-behaviors, the ‘epoch’ model (where the periodicity depends only on the sleep epoch) emerged as the winning model and a reliable main effect of epoch was found (p<0.001) in all cases. Further, we performed post-hoc tests using tukey adjustment (for multiple comparisons) to identify differences between pairs that are significant. Thus, we found an apparent increase in the likelihood of periodicity for both antennae and proboscis during the middle segments of sleep bouts (*Figure 1F,G*). This suggested physiological differences which might be detected in the fly brain, so we then performed electrophysiological recordings in a similar context.

### Long-term multichannel recordings with spontaneous sleep bouts

We recorded local field potentials (LFPs) across the fly brain using a linear 16-channel electrode inserted into the left eye of flies in a similar context as above, walking (or resting) on an air-supported ball (*Figure 2A,B*). The electrode insertion location was positioned to sample LFPs from the retina to the central brain (Paulk et al. 2013) (*Figure 2C*, white arrowheads). The depth of insertion of the electrode was optimized by using a visual stimulus calibration protocol, based on a reliable LFP polarity-reversal identified in the fly inner optic lobes (*Figure S2*; and see Methods for polarity reversal). The change in polarity (positive to negative deflections in response to the visual stimulus) was always positioned between electrodes 11-13 in all flies, before the start of the long-term LFP recordings. This LFP polarity-based method allowed us to maintain a level of recording consistency across flies in terms of spatial locations of the electrodes, thereby allowing us to compare and combine LFP data across multiple flies. To further ensure reproducible recording locations, we also developed a dye-based registration method (*Figure S3,4*; and see Methods for dye-based localization) and estimated recording channel locations in the brain for two sample flies. Using this method we identified three broadly-defined brain recording regions to simplify our subsequent analyses (*Figure 2D*): central channels (1-5), middle channels (6-10) and peripheral channels (12-16); here assuming polarity reversal in channel 11. Also for further analysis, as the polarity reversal channel is used for re-referencing, the number of channels used in analysis becomes 15.

**Figure 2:**
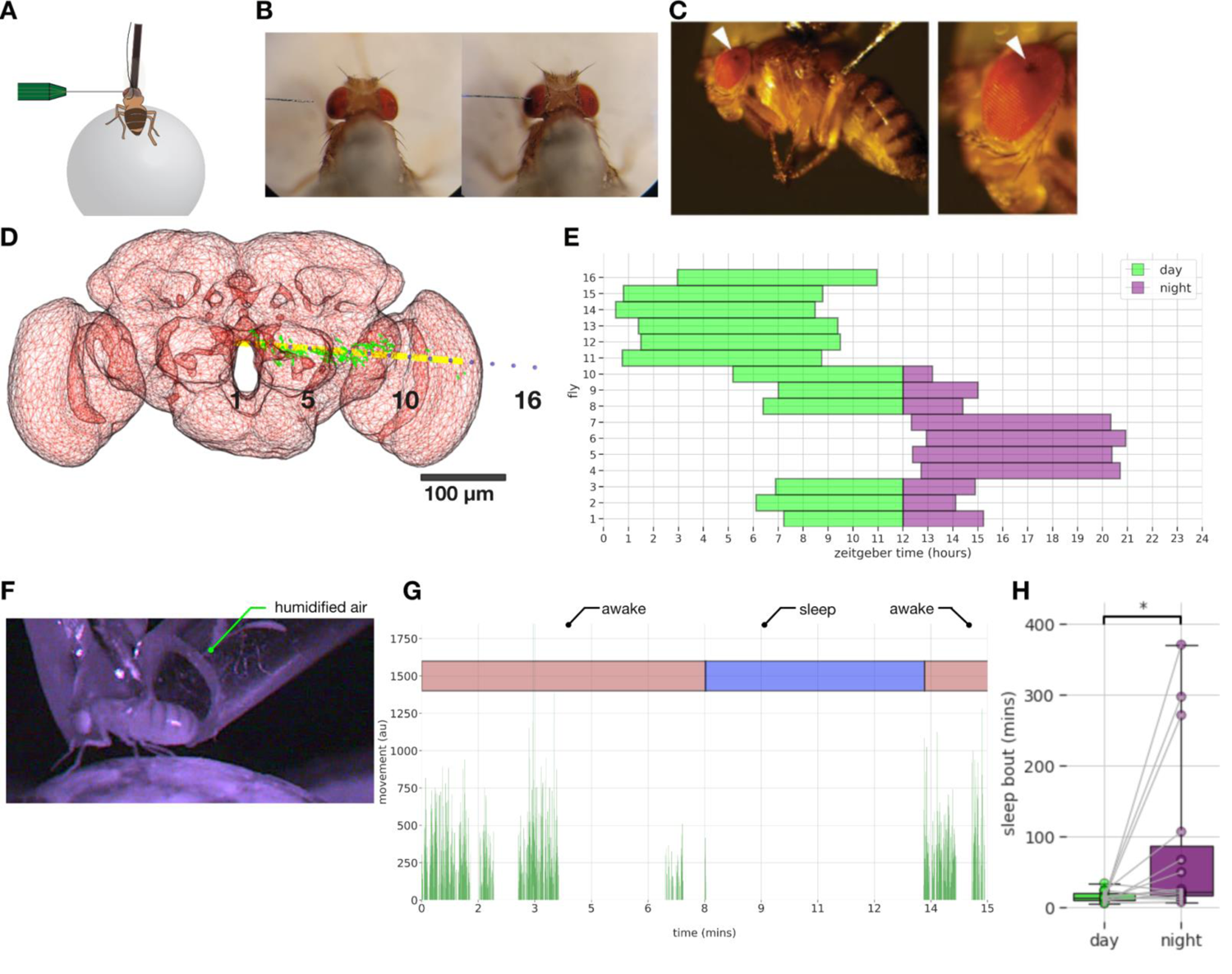
A) In vivo long-term electrophysiology recording setup: tethered flies were placed on an air supported ball setup which served as a platform for walking/rest. B) Top view of electrode insertion process, with electrode approaching from the left eye of an example fly. C) Side view of electrode insertion site on the dorsal part of the left eye. D) Localization of electrodes using fluorescent dye: the electrode numbers (black) are displayed along with the dye (green), eigenvector (yellow) indicating the main path of the probe, the fafb14 neuropil (red). E) Raster plot showing the used recording times (LFP) for the 16 flies. Only the first 8 hours of LFP recording were used for analysis though all the flies survived for more than 24 hours. F) Fly movement is quantified using video recorded in profile view with infrared lighting. Movement was quantified between adjacent frames with pixel difference and contour thresholding. G) Movement area (activity pattern) plotted along with ‘awake’ and ‘sleep’ state labeling for an example sleep bout. H) Sleep bouts during day are significantly longer than night thus confirming the occurrence of natural sleep in our setup. * indicates p < 0.05

We utilized the above calibration steps and recorded LFP data from 16 flies over the course of a day and night cycle (*Figure 2E*; and See Methods for data exclusion criteria). We designed our recordings so that experiments were started at different times in different flies, to achieve complete coverage of a full day-night cycle. We however only examined the first 8 hours of the LFP data in each fly (*Figure 2E*), to ensure we were always recording from active and responsive animals (all 16 flies were still alive after 24 hours).

The behavior of the flies was recorded under infrared lighting (*Figure 2F*) and their movements were quantified using a combination of pixel difference (van Alphen et al. 2013) and contour thresholding between neighboring frames (See Methods for movement analysis). Sleep was defined by 5 min immobility criteria, based on previous observations in unrestrained flies (P. J. Shaw et al. 2000; Hendricks et al. 2000) as well as tethered flies (van Alphen et al. 2013; Yap et al. 2017). Fly mobility along with classification of different behavioral states (‘awake’, ‘sleep’) for an example sleep bout is shown in *Figure 2G*. Since it was unclear whether flies would even sleep in this multichannel recording preparation, we tallied immobility bout durations across the day and the night for each fly (we used 16 hrs of video data for each fly - See Methods for data exclusion criteria), expecting that flies should be sleeping more at night on average. We found that flies were able to sleep in this preparation, and that nighttime sleep bouts were indeed longer than daytime sleep bouts (median = 22.42 min vs 13.99 min, respectively; t(13) = −2.32, p<0.05) (*Figure 2H*). This confirms that similar to single channel LFP recordings (van Alphen et al. 2013; Yap et al. 2017) flies slept reliably in this multichannel recording preparation, allowing us to assess changes in LFP activity across the fly brain during sleep and wakefulness, and to relate these changes to sleep micro-behaviors.

### LFP differences across the brain during spontaneous sleep and awake

Next, we focused on the multichannel data to identify potential differences between sleep and wake across the fly brain, separating our recordings into three broad regions: central, middle, and peripheral *(Figure 3A)*. An example sleep bout and its corresponding spectrograms across the central, middle, and peripheral channels reveals increased activity during sleep in the central brain compared to the periphery *(Figure 3B)*. Additionally, we noted variegated effects in the lower frequencies (5-10 Hz) within the sleep bout *(Figure 3B,* arrowheads) as well as significant LFP activity (5-40 Hz) associated with locomotion.

**Figure 3:**
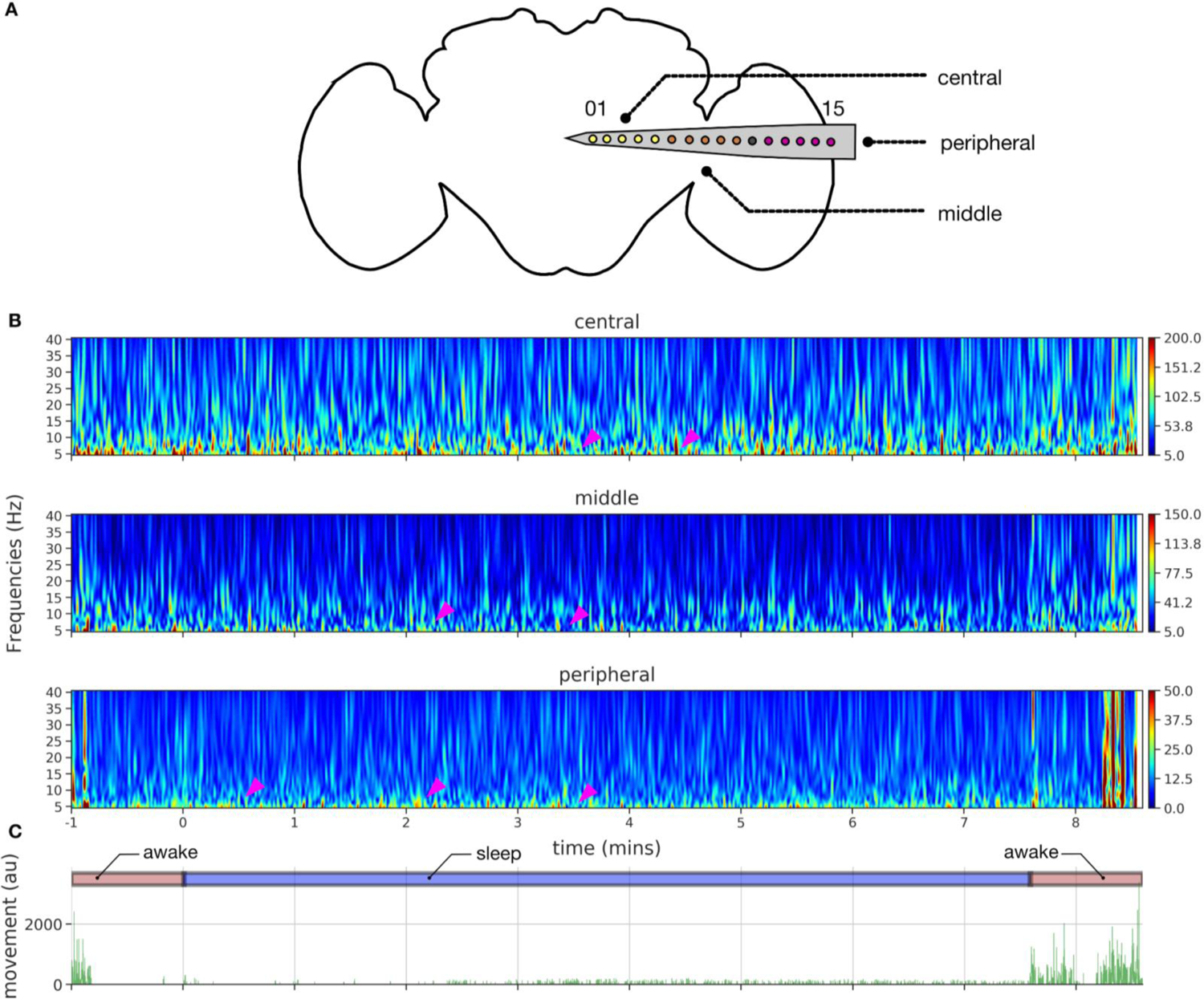
A) 16 electrodes color coded by location (purple - peripheral, gray - reference, brown - middle, yellow - central) are illustrated on an outline of a standard *drosophila* brain. B) Spectrogram in different channels groups across an example sleep bout shows variation (magenta arrowheads) in the lower frequency bands (5-10 Hz) within the sleep bout, while activity across 5-40 Hz in the flanking awake period. C) Movement area (activity pattern) plotted along with ‘awake’ and ‘sleep’ state labeling for this example sleep bout.

When we examined sample LFP data more closely across all channels (*Figure 4*), we observed higher LPF amplitudes in the central and middle channels than in the peripheral channels, and more activity during wake than during sleep (*Figure 4A,B*).

**Figure 4:**
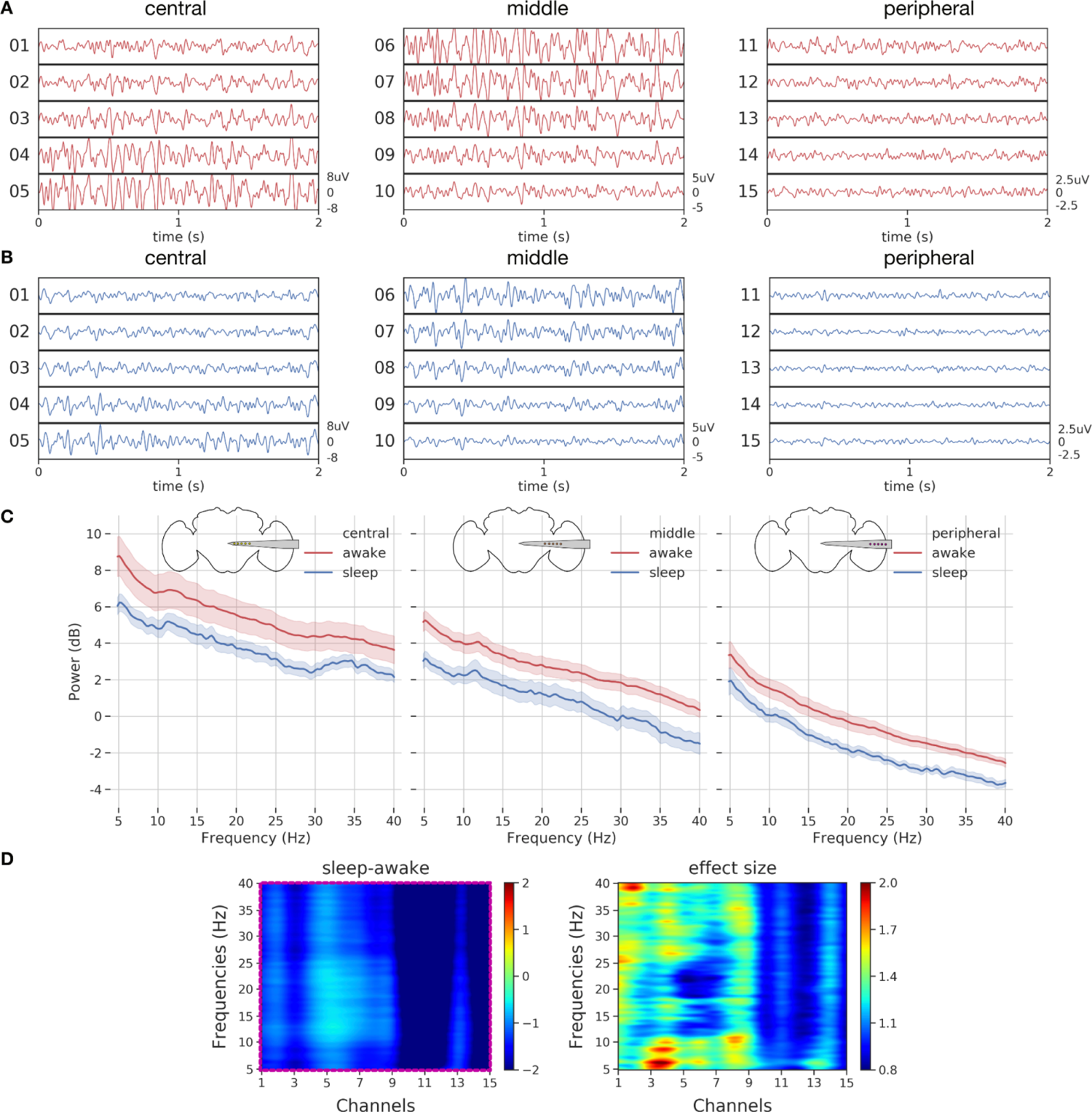
A,B) Average LFP across a sample awake and sleep bout in an example fly. C) Mean power spectrum of LFP (5-40 Hz) across ‘awake’ and ‘sleep’ states in the central, middle, peripheral channels. Across all channels, ‘sleep’ periods have lower LFP power compared to the ‘awake’ periods. D) Spectrogram showing the mean difference across ‘sleep’ and ‘awake’ periods, while clustering analysis reveals a single significant cluster (magenta box) across all channels and frequencies. Effect sizes are also plotted to identify the individual effect values for every frequency and channel pair.

Interestingly, the fly brain is not necessarily quiet during sleep, with some channels (e.g., channels 5-7) displaying increased activity compared to other channels. To substantiate our observations, we performed spectral analysis on the data. For this purpose, we epoched the LFP data into 60 sec bins and computed the power spectrum per epoch per channel (See Methods for LFP analysis - preprocessing, power spectrum analysis). Since LFP data recorded from flies can be sensitive to physiological artifacts such as heartbeat and body movements (Paulk et al. 2013), we employed a common referencing system (based on a brain based signal) that allowed for removal of non-brain based physiological noise. Plotting the power spectral density across the three different channel groupings for different frequency bands (5-40 Hz), revealed consistently greater power in all flies (n=16) during wake than during sleep across the entire recording transect (*Figure 4D*). Although decreased LFP power during sleep is consistent with previous findings involving single channel recordings in flies (van Alphen et al. 2013; Yap et al. 2017; Nitz et al. 2002), it was surprising to see that even the fly optic lobes are significantly less active during sleep compared to wake, suggesting a brain-wide effect.

We next examined more closely the relationship between individual channels and LFP spectral frequency between sleep and wake states. We employed non-parametric resampling tools to identify the precise patterns (frequency x channel pairs) differing across awake and sleep at the group level. For this purpose, we first computed the difference in mean spectral data across awake and sleep for individual flies. Then, we performed a cluster permutation test (flies x frequencies x channels) on the difference between awake and sleep data (*Figure 4D - left panel*) to reveal a significant cluster (frequency x channel pair). The significant cluster as indicated by the magenta box (*Figure 4D - left panel*) covered all frequencies (5-40 Hz) and channels (1-15), thereby confirming the spectral results in *Figure 4C* that found a brain-wide decrease in power during sleep compared to wake. As we had a single significant cluster (magenta box), we then sought to identify subclasses of frequencies and channels within this cluster which might be more specifically associated with sleep.

We computed the effect sizes for every channel x frequency combination (*Figure 4E - right panel*). This revealed an interesting frequency structure distinguishing sleep from wake. This included areas of interest in the 5-10 Hz and 25-40 Hz range in the central channels (1-3). The 5-10 Hz frequency domain was identified in a previous study as being relevant to sleep in *Drosophila* (Yap et al. 2017), and the higher 25-40 Hz range overlaps with the frequencies associated with attention-like behavior in flies (Bruno van Swinderen and Greenspan 2003; Grabowska et al. 2020). Consistent with previous work, it is however clear that LFP activity is mostly decreased during all of sleep compared to wake, even in the 7-10 Hz range that has been associated with certain sleep stages (*Figure S5*).

### LFP differences across induced sleep and awake

Sleep can be acutely induced in *Drosophila* by using optogenetic or thermogenetic activation of sleep-promoting neurons (Shafer and Keene 2021). We were curious whether induced sleep revealed similar effects across the fly brain, following the same statistical approaches employed above for spontaneous sleep. For this, we focused on whole-brain recordings taken from 104y-Gal4 / UAS-TrpA1 flies, a sleep-promoting circuit (*Figure S6A*) that expresses a temperature sensitive cation channel in the fan-shaped body in the central brain (Donlea et al. 2011). As shown in a previous study (Yap et al. 2017) as well as other *Drosophila* sleep studies (Dag et al. 2019), activating these neurons with heat (temperature ∼ 29°C) results in behavioral quiescence and induced sleep, whereas control strains remain awake and active. In these recordings, a different multichannel probe was employed (*Figure S6B*), with 16 recording sites that spanned the entire brain from eye to eye (Paulk et al. 2013). We preprocessed the induced sleep LFP data (See Methods for thermogenetic sleep induction) in a similar fashion to the spontaneous sleep LFP data. We first contrasted the mean power spectra per fly under two conditions: baseline and sleep induction (*Figure S6C*). As above, we then performed a cluster permutation test (flies x frequencies x channels) on the difference between baseline wakefulness and induced sleep, to reveal a significant cluster (frequency x channel pair). Thus, we uncovered a significant cluster (*Figure S6D*) in the central brain channels across all (5-40Hz) frequency bands, whereas the 104y-Gal4/+ control flies did not reveal such a cluster (*Figure S6E,F*). It is interesting to note that sleep induction using this strain yielded an opposite effect to what we found during spontaneous sleep: LFP activity during induced sleep is on average higher than during baseline wakefulness (*Figure S6D*), while it was lower during spontaneous sleep (*Figure S5*). Additionally, the effect observed during induced sleep was only observed in the central channels whereas the spontaneous sleep effects appear to at least cover the entire hemisphere from center to periphery. This shows that genetically-induced sleep in flies can produce strikingly different electrophysiological signatures than spontaneous sleep, consistent with several previous similar observations (Tainton-Heap et al. 2021; Yap et al. 2017; Anthoney et al. 2023; Lin, Panula, and Passani 2015; Troup et al. 2018). For the rest of this current study, we focus on spontaneous sleep.

### Machine learning identifies distinct sleep stages in multichannel data

Our earlier analysis of micro-behaviors during sleep in this preparation (*Figure 1*) suggests that sleep is not a single phenomenon, and that the requisite 5 min immobility criterion might not fully capture potential LFP and behavioral changes that might occur across a sleep bout. There is evidence that sleep quality (via arousal threshold probing) in wild-type *Drosophila* flies also changes across a bout of quiescence (van Alphen et al. 2013; Faville et al. 2015), suggesting that flies transition from lighter to deeper sleep stages. To assess whether this might also be evident in our multichannel recordings, we divided our LFP data (for all channels) into five different temporal segments, analyzing only sleep epochs that were 5 min or longer (*Figure 5A*): 1) ‘presleep’: the 2 mins (−2 to 0 mins) before flies stopped moving; 2) ‘earlysleep’: the first 2 mins (0 to 2 mins) after the start of a sleep bout; 3) ‘latesleep’: the last 2 mins of sleep before mobility resumed; 4) ‘midsleep’: any time between ‘earlysleep’ and ‘latesleep’. 5) ‘awake’: the rest of our LFP data. Our partitioning of the LFP data matches a similar partitioning applied to whole-brain calcium imaging of flies engaged in spontaneous sleep (Tainton-Heap et al. 2021).

**Figure 5:**
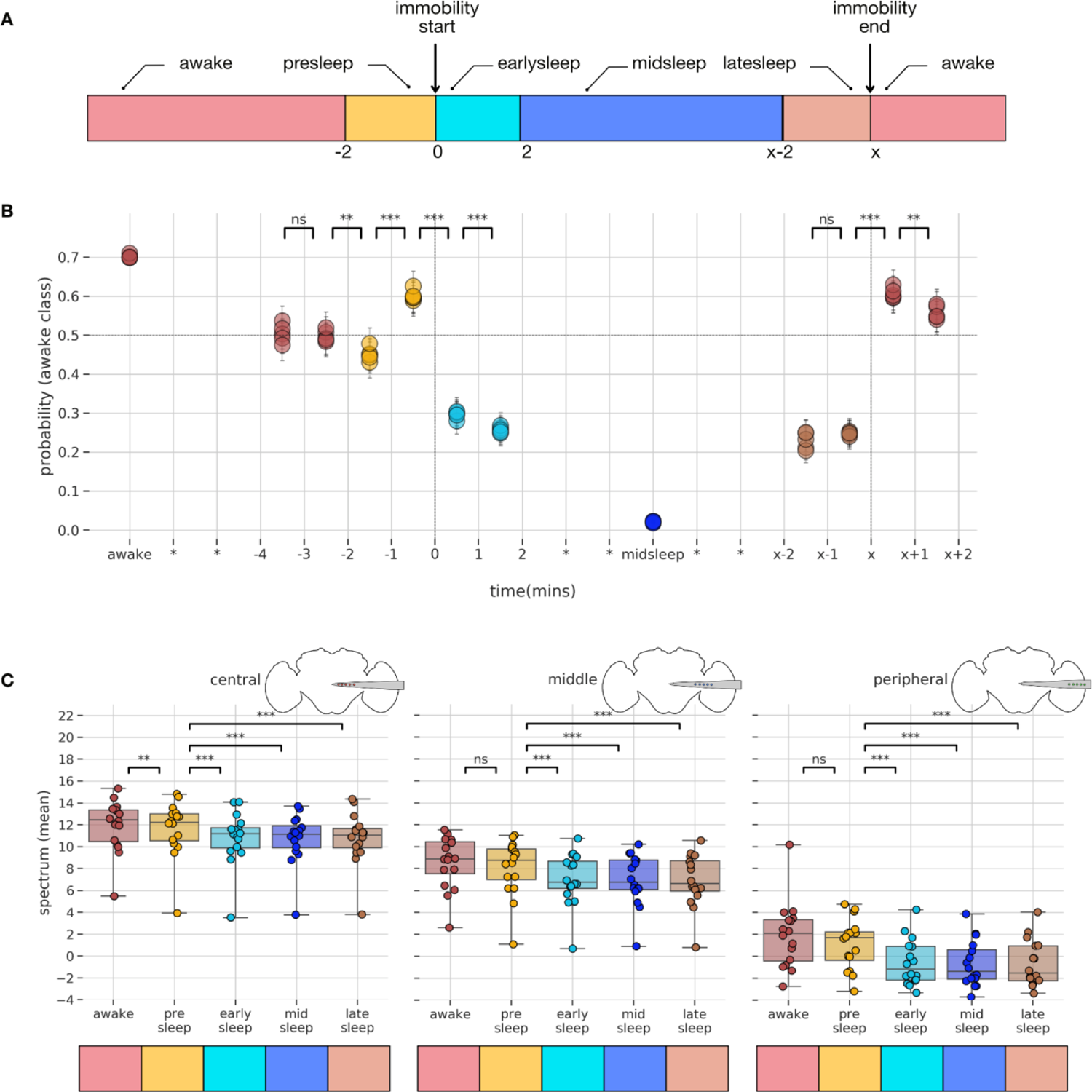
A) Sleep bouts (> 5 min) were binned into 4 segments. 2 mins before the start of immobility (presleep), 2 mins after the start of immobility (earlysleep), 2 mins before the end of immobility (latesleep), the period between early and latesleep is midsleep, rest of the periods are categorized as awake. B) Probability estimates (of awake class) plotted across different time segments. Horizontal dotted line indicates values above 0.5 are likely to be classified as awake and below 0.5 as sleep. Vertical dotted line at 0 indicates start of immobility period and at x indicates end of immobility period. Unpaired samples t-test was conducted across different epochs to test for statistical significance. C) Comparison of mean power spectrum across different channels and different sleep stages. ** p<0.01, ***p<0.001, ns indicates not significant.

To understand how LFP based signatures change within a sleep bout, we decided to perform a hypothesis-agnostic analysis through machine learning techniques. To perform such machine learning based classification, we first used support vector machine (SVM) based techniques. Briefly, SVM belong to a class of supervised learning model, that is comprised of building a hyperplane or set of hyperplanes in a high dimensional space (using the kernel trick for non-linear mapping functions) with the goal to maximize the separation distance between the closest data point (in the training dataset) of any class (functional margin) (Cortes and Vapnik 1995). The choice of the optimal hyperplane is made in such a way that the generalization error would be lower for the new data points in the test dataset (*Figure S7A*). For detailed steps for preprocessing of data and implementation of classifiers refer to Methods for sleep staging by classifiers. The probabilistic prediction per class per iteration is shown in *Figure 5B*. It is interesting to note several points. First, the probability of awake data is ∼0.7 and of midsleep is ∼0.0 indicating that the classifier performs well on classes that it has already been trained on. Second, at the epoch −2 to −1min, when the fly is still moving (yellow circles), LFP data indicates that it is closer to resembling sleep (<0.5), before dropping fast to ∼0.3 (turquoise circles) in the first two minutes of sleep.

The above analysis indicates that with this approach we could predict the probability a fly will fall asleep 2 mins before the start of the immobility period. Interestingly, just 1 min before flies fall asleep the LFP data indicates a brief moment more closely resembling wake (yellow circles), perhaps associated with grooming periods (observed in honeybees for example (Eban-Rothschild and Bloch 2008)). Interestingly, in the first two minutes of sleep (turquoise circles) reveal a probability metric halfway between midsleep and wake, suggesting either a gradual descent into deeper sleep or a distinct sleep stage. Finally, at the epoch from x-2 to x-1 min before mobility resumes (brown circles), the probability metric returns to a similar level as early sleep. Immediately after mobility resumes, the LFP data is classified as no different than awake, i.e, there is no post-sleep ambiguity. It is important to note that only the ‘awake’ and ‘midsleep’ data has been seen by the classifier, the rest of the data −4 to +2 min, x-2 to x+2 min has never been seen by the classifier. Additionally, midsleep collapses a wide range of different sleep durations in different flies, so could still be averaging different sleep states within. Nevertheless, our results suggest that broadly dichotomizing mid sleep and wake identifies other sleep stages that resemble neither.

### Model based spectral analysis differentiates wakefulness from sleep bouts across different channels

Having revealed how multichannel LFP data can be used to differentiate across different temporal stages of sleep, we next decided to identify what channels might be important for revealing this. For this purpose, we employed a multilevel modeling approach. To reveal how spectral data might change throughout the fly brain across a sleep bout, we calculated the mean spectral power for each of the aforementioned epochs and pooled data from central, middle, and peripheral channels. Because different flies had varying numbers of sleep epochs, we used multilevel models instead of traditional repeated measures of analysis of variance. For details refer to the methods section (Multilevel models - models for spectral analysis). The ‘epoch-channel’ model emerged as the winning model; here the power spectrum depends on a combination of the LFP epoch type and the channel type. In the epoch-channel model, we found that there was a reliable main effect of both epoch (p<0.001) and channel (p<0.001) on power spectrum and also the interaction between epoch and channel also had a reliable effect (p<0.001) on power spectrum. In summary, the above model-based analysis confirms that the power spectrum of the LFP data varies based on the channel location and also the epoch state of the fly.

We then proceeded to examine more closely how differences in the sleep LFP might be segregated across the fly brain (*Figure 5C*) using post-hoc tests (using tukey adjustment for multiple comparisons) from the epoch-channel model. In the central channels, the ‘awake’ data was significantly different compared to all sleep categories, and critically was also different to the ‘presleep’ data. It is important to note that behaviourally the fly is still considered awake in the ‘presleep’ period (i.e., it is still moving). Thus, the ability to predict sleep at least 2 mins before the onset of immobility, which was revealed in our SVM analysis (*Figure 5B*), might be explained by these significant spectral differences only observed in the central channels. In the middle channels, the ‘awake’ data was also significantly different across all sleep categories, however was not different to the ‘presleep’ data. Further, the ‘presleep’ period was significantly different from ‘earlysleep’,‘midsleep’,‘latesleep’ periods. In the peripheral channels, the ‘awake’ data was significantly different across all sleep categories, however was again not different to the ‘presleep’ data. Taken together, mean power spectral data across different channels was thus able to differentiate between ‘awake’, ‘presleep’, and different sleep epochs of sleep. However, the post-hoc analysis did not differentiate among sleep epochs (‘earlysleep’, ‘midsleep’, ‘latesleep’). Since this is inconsistent with previous findings using single glass electrodes (Yap et al. 2017), we questioned if the pooling of channel x frequencies data (3 broad brain regions x 1 overall power spectrum) could be hiding more specific effects which might become evident with the full (15×145) dimension of channels x frequencies.

### LFP features across different temporal stages of sleep

Having established the existence of different temporal stages of sleep using a classifier based on SVM and confirming the same using model-based analysis, we were next interested in the features in the LFP data (which channels at what frequencies are important for distinguishing epochs within a sleep bout), that helps us differentiate these stages.

For this purpose we used random forest classifiers. A random forest classifier is a class of supervised learning algorithms that utilizes an ensemble of multiple decision trees for classification/regression. This could be illustrated by an example (*Figure 6A*). In the first step subsets of training data (#1 to #n) were created by making a random sample of size N with replacement. This allows for the ensemble of decision trees (#1 to #n) to be decorrelated and the process of such random sampling is called bagging (bootstrap aggregation). In the second step, each decision tree (#1 to #n) picks only a random subsample of features (feature randomness) instead of all features (again allowing for the decision trees to be decorrelated).

**Figure 6:**
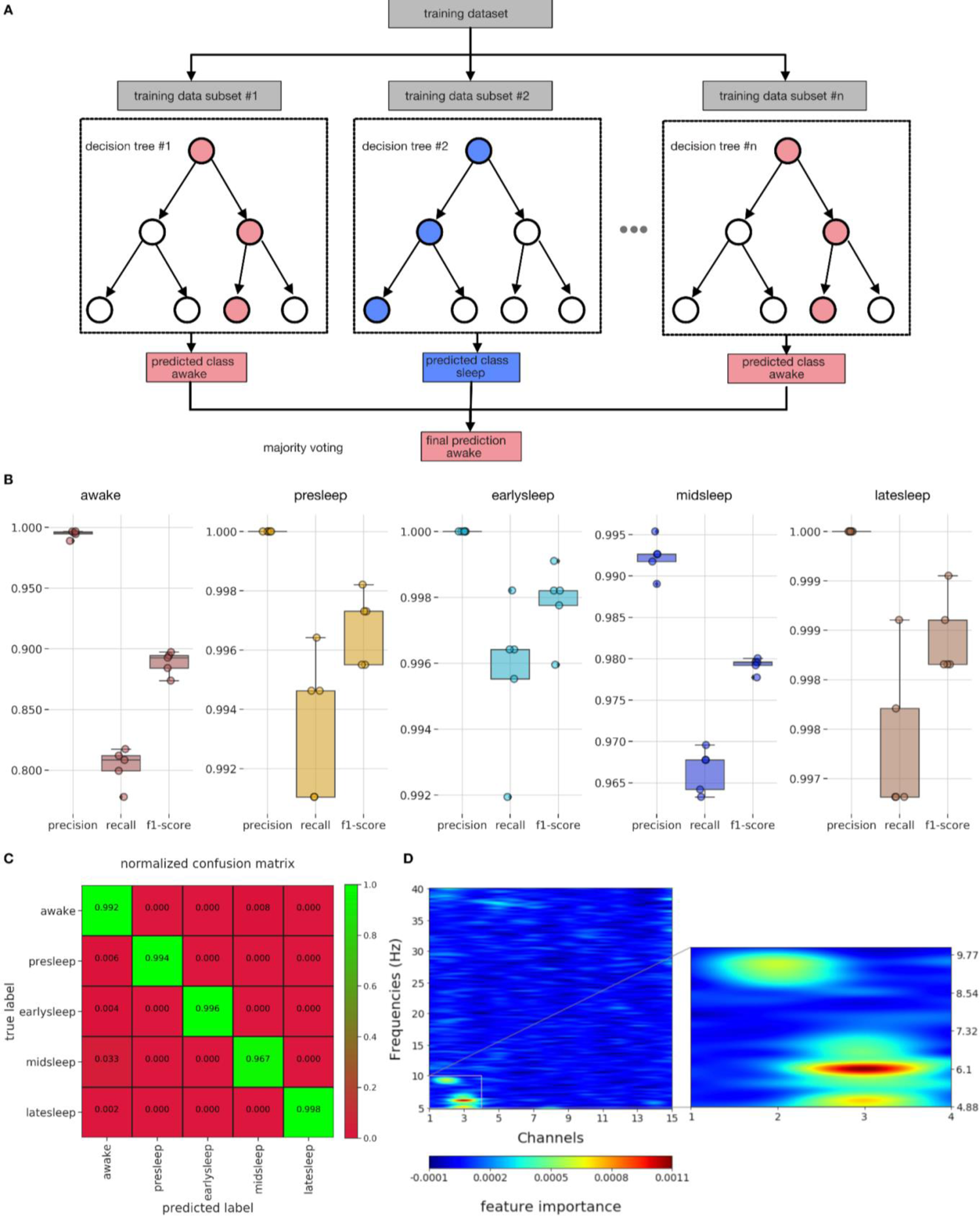
A) Schematic indicating the workings of a multiclass random forest classifier in identifying the predicted class. B) Performance metrics like precision, recall, f1-score across different classes of the trained classifier. C) Normalized confusion matrix of the trained classifier. D) Feature importance of the multiclass classifier indicates an ROI across central channels and frequency bands (5-10 Hz) as critically important.

In the final step, all the decision trees create individual predictions of classes and the final outcome would be resolved by simple majority voting (illustrated here with a goal of classifying ‘awake’ vs ‘sleep’). Thus, bagging and feature randomness allows for the random forest to perform better than individual decision trees.

We performed a multiclass classification of the following classes: ‘awake’, ‘presleep’, ‘earlysleep’, ‘midsleep’, ‘latesleep’. For the detailed preprocessing and feature computation steps refer to the methods section (Sleep staging by classifiers - multiclass svm analysis & feature importance). We then computed classifier performance metrics (See methods for sleep staging by classifiers - classifier metrics) like precision, recall, f1-score (*Figure 6B*) and further normalized confusion matrix (*Figure 6C*) which reveal excellent performance in predicting the multiple classes (green boxes). This indicates that classifier features (channels x frequency) are sufficient to distinguish multiple sleep stages (classes) and furthermore provide direct evidence of multiple sleep stages. Another reason for using random forest classifiers is that it is possible to identify relative feature importance in the performance of classifiers, thereby identifying features (channels x frequency) which are important for differentiating across multiple sleep stages.

To identify the LFP features most likely discriminating among sleep stages, we utilized the multiclass random forest classifier (described above), and uncovered the features that are important in this classifier (*Figure 6D*) with permutation importance technique. Interestingly, the most important features fall within a narrow range of channels (1-3) and frequencies (5-10 Hz). This indicates that the 5-10 Hz frequency range within the central channels are the most important in resolving different sleep stages. Next, we decided to cross-validate the utility of this permutation-based technique in resolving across different epochs. For this purpose, we created a multiclass random forest classifier, with target classes as: ‘awake’, ‘sleep’, and identified the features that are important in this classifier (*Figure S8A*). The most important features are actually distributed evenly among all the features (channels x frequency), thus cross-validating our previous clustering results (*Figure 4D*) wherein we showed that the LFP differences across ‘awake’ and ‘sleep’ are distributed across all channels and frequencies.

### Proboscis extension behavior during sleep in multichannel recordings

Earlier, we identified rhythmic proboscis extensions (PEs) during midsleep (*Figure 1*), which we propose describe a distinct sleep stage in *Drosophila* (van Alphen et al. 2021). However, it is unclear if brain activity associated with PEs are sleep-like or PE-specific. This distinction is important, as it would disambiguate a unique brain state (deep sleep) from a specific behavior associated with that state (PEs). In order to identify PEs in our electrophysiological dataset, we again used DeepLabCut (Mathis et al. 2018) to track different body parts of the fly (*Figure 7A*). We further used multiple classifiers based on the tracking data, followed by manual verification to identify the PEs. Sample proboscis extension periods in an example fly along with a few of the features (x,y proboscis location, likelihood of location, distance of proboscis to eye) are shown in *Figure 7B*. For more details on the proboscis detection steps refer to the section - Methods for proboscis tracking for flies on electrophysiology setup. Our classifier accuracy was over 80% for most flies (*Figure 7C*): the ground truth was validation by a human observer on classifier detected events. In *Figure 7D*, we plot the mean proboscis to eye distance for all the flies averaged across awake and sleep bouts. As described earlier for flies without implanted electrodes, PEs executed during wake and sleep are behaviourally similar and hence would be difficult to distinguish from each other using video alone. Similar to our behavioral dataset, PE events usually occur in rhythmic bouts of more than one, rather than single events. In *Figure 7E*, we plot the inter-proboscis interval period, which is the interval between consecutive PE events in a single proboscis bout. It can be seen that most proboscis events occur within 1.8 sec (95^th^ percentile) of each other. As shown before in our behavioral data without implanted electrodes, the inter-proboscis interval does not vary across awake and sleep periods. Next in *Figure 7F*, we decided to probe the number of single (one PE event) and multi (>1 PE event) across different flies. We found that occurrences of single PE events are significantly lower than multi PE events using a pairwise t-test with t(13) = 3.72, p<0.01.

**Figure 7:**
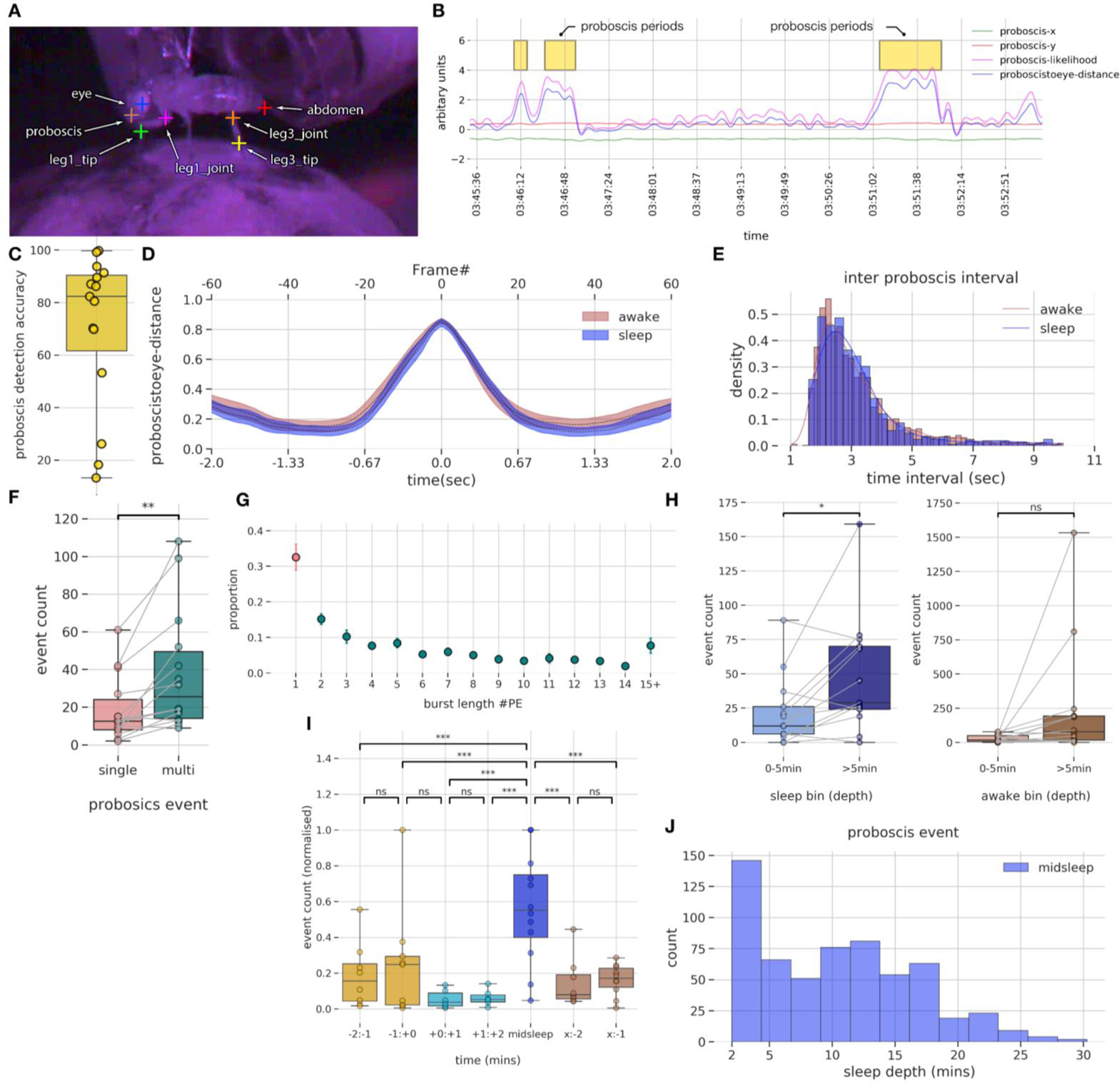
A) Seven different body parts were annotated using *DeepLabCut* for pose estimation. B) Identified PE periods (yellow boxes) were plotted along with filtered body parts of proboscis and other estimated metrics. C) PE events detection accuracy across different flies. D) Average proboscis to eye distance (across all flies) plotted across frames and time periods are similar in awake and sleep states. E) During PE burst events, inter proboscis intervals are highly regular, with one PE occurring every 1.5 s at group level. F) PE events are more likely to occur as multiple events (bursts) instead of a single event. G) About 33% of PE events occur as single while the rest are bursts of varying length. H) Number of PE events occurring after the first 5 mins of sleep is significantly higher than in the first 5 mins, indicating that more PE events occur in deeper stages of sleep. Also displayed is the control analysis, with awake depth showing no increase with PE event count. I) Normalized proboscis event count across different sleep segments: −2:−1 indicates 2 mins before start of sleep, −1:0 indicates 1 mins before start of sleep, +0:+1 indicates 1 mins after start of sleep, +1:+2 indicates 2 mins after start of sleep, x:-2 indicates 2 mins before end of sleep, x:-1 indicates 1 mins before end of sleep. The normalized count is significantly higher in the midsleep segments compared to other segments. J) Proboscis events occurring in midsleep across different sleep depths. *p<0.05, ** p<0.01, ***p<0.001, ns indicates not significant.

To further illustrate this point in *Figure 7G*, we plotted the burst length of a PE event (number of extension events within a PE bout) and found that only 33% of the events are single PE while the rest are multiple PE events. Overall, our investigation of PEs in this multichannel recording dataset is in concurrence with our first (electrode-free) dataset, suggesting that inserting probe into the fly brain does not alter several measures associated with this micro-behavior.

Previous work has linked PEs with a deep sleep stage in flies (van Alphen et al. 2021). We therefore next investigated whether the number of PEs varied across a sleep bout in our LFP recording dataset, as suggested in our purely behavioral dataset (*Figure 1G*). We found that more PE events occur after 5 min of a sleep bout, compared to those occurring before the 5th min of a sleep (*Figure 7H*) (pairwise t-test, t(12) = −2.8, p<0.05), suggesting that PEs indeed predominate during deeper sleep. We also compared PEs immediately after flies had awakened from sleep, which revealed no significant difference (*Figure 7H*) (pairwise t-test, t(13) = −1.92, p>0.05) between PE bouts occurring after the 5th min of an awake bout compared to those occurring before the 5th min of an awake bout, confirming that transitions into sleep (rather than transitions back to wake) were associated with increased PE events.

We next asked if the number of PE events changed across a sleep bout in our multichannel recording preparation. To determine if the PE event count varies across different temporal sleep stages (*Figure 7I*) we used multilevel models. For details refer to the methods section (Multilevel models - models for PE event counts). The time_label model (where the PE event count depends only on the specific temporal sleep stage) emerged as the winning model. Further, we performed post-hoc tests using tukey adjustment (for multiple comparisons) to identify differences between pairs that are significant. We found that PE events occur more often in midsleep compared to other sleep stages. Returning to our original observation that most PEs occur after 5min of sleep, we plotted the distribution of PE events occur in the midsleep epoch across all flies (*Figure 7J*), and found that 95 percentile of all PE events in midsleep indeed occur after 2.5 minutes of the midsleep epoch (thus, 4.5 mins from sleep onset).

### LFP features of a deep sleep stage with proboscis extension

We next questioned whether PEs occurring during sleep and wake had similar neural correlates, or if the sleep-related events were indeed different and thus indicative of a unique sleep-related function. We therefore focused on the multichannel data to identify any differences in the LFP activity associated with PEs during wake and sleep epochs. We first identified the PE periods (Refer to Methods LFP analysis - proboscis: Identification of proboscis periods) and extracted the LFP data and epoched them into 1 sec bins. Second, we used spectral analysis to determine if epochs characterized by PEs differ in frequencies across different channels, for wake compared to sleep. For this purpose, we computed the spectral power for every 1 sec epoch per channel (See Methods for LFP analysis - proboscis: power spectrum analysis), using as before a common reference system for re-referencing the LPF data. Third, we employed non-parametric resampling tools to identify the precise patterns (frequency x channel pairs) differing in proboscis periods within awake and sleep at the group level. For this purpose, we first computed the difference in mean spectral data across non-proboscis periods (awake or sleep) and proboscis periods (awake proboscis and sleep proboscis respectively) for individual flies. We then performed a cluster permutation test (flies x frequencies x channels) on the difference data to reveal significant clusters (frequency x channel pair).

In *Figure 8A*, we show the difference data (awake proboscis - awake period) and clustering analysis, which reveals a significant cluster in the middle channels (6-10) across all frequencies. Further, within the significant cluster we also performed a post hoc analysis revealing that spectral activity within the awake proboscis periods are lower than awake periods. In *Figure 8B*, we show the difference data (sleep proboscis - sleep period) and clustering analysis reveals a significant cluster in the central channels (1-5) across higher frequencies (32-40 Hz). Further, within the significant cluster we also performed a post hoc analysis revealing that spectral activity within the sleep proboscis periods are higher than sleep periods (in contrast to the awake proboscis periods). In *Figure 8C*, we directly compared the awake and sleep proboscis periods and showed the difference data (awake proboscis - sleep proboscis) and clustering analysis, which reveals a significant cluster in the central, middle channels (1-9) across higher frequencies (25-40 Hz). Further, within the significant cluster we also performed a post hoc analysis revealing that spectral activity within the sleep proboscis periods are lower than awake proboscis periods. This suggests that PEs occurring during sleep are qualitatively different from identical PE events occurring during wake. This suggests that the brain activity state (e.g., quiet or deep sleep (Tainton-Heap et al. 2021; Anthoney et al. 2023)) overrides the neural correlates associated with the same behavior occurring during wake.

**Figure 8:**
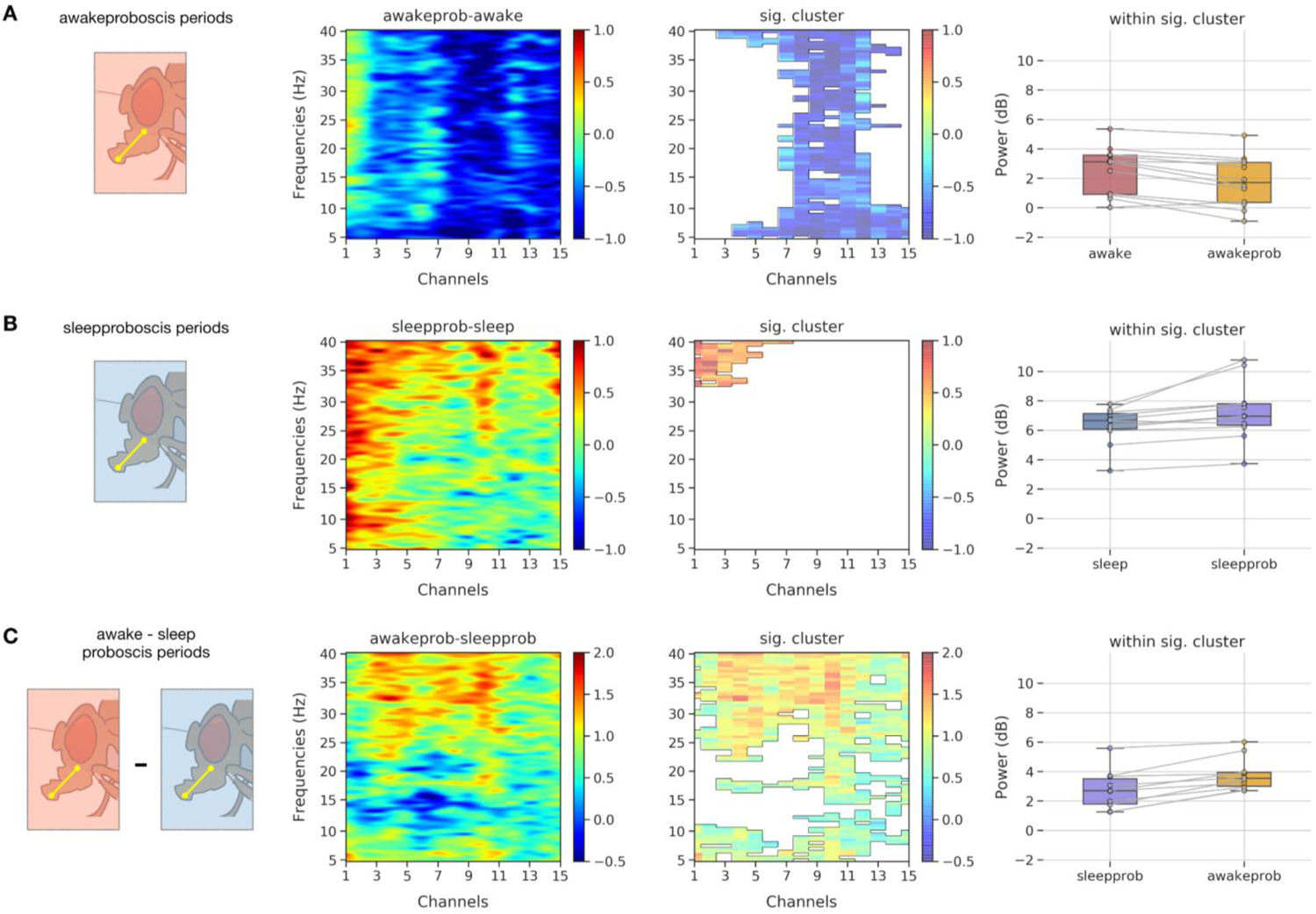
A) Spectrogram showing the mean difference across ‘awakeprob’ (PE events in awake periods) and ‘awake’ periods, while clustering analysis reveals a single significant cluster across middle channels in all frequencies. Activity within the significant cluster indicates activity in the ‘awakeprob’ is comparatively lower than ‘awake’ periods. B) Spectrogram showing the mean difference across ‘sleepprob’ (PE events in sleep periods) and ‘sleep’ periods, while clustering analysis reveals a single significant cluster across central channels in higher frequencies (35 - 40 Hz). Activity within the significant cluster indicates activity in the ‘sleepprob’ is comparatively higher than ‘sleep’ periods. C) Spectrogram showing the mean difference across ‘awakeprob’ and ‘sleepprob’ periods, while clustering analysis reveals a single significant cluster mostly across all channels in higher frequencies (25 - 40 Hz). Activity within the significant cluster indicates activity in the ‘sleepprob’ is comparatively lower than ‘awakeprob’ periods, thereby elucidating a significant difference across proboscis events occurring in sleep and awake periods (though phenotypically they look the same - Figure 7D).

## Discussion

Sleep is most likely a whole-brain phenomenon, meaning that its presumed varied functions (Kirszenblat and van Swinderen 2015) are understood to be of benefit to the entire brain rather than to only specific sub-circuits. There is good evidence for this in the *Drosophila* model, with synaptic physiology for example changing during sleep in the optic lobes of flies (Donlea, Ramanan, and Shaw 2009) as well as brain-wide (Gilestro, Tononi, and Cirelli 2009). Similarly, in mammals, subcortical as well as cortical brain regions experience sleep-related changes that are thought to be important for maintaining neuronal homeostasis (Tononi and Cirelli 2014). Accordingly, to better understand sleep in an animal model such as *Drosophila melanogaster* requires sampling associated changes in neural activity across the fly brain, and not only in specific sub-circuits of interest. Unlike in larger animal models such as mice, recording from multiple brain regions in behaving (and sleeping) flies has been challenging, so there has been limited capacity to investigate dynamic brain processes during sleep in this otherwise powerful model system. While genetically encoded reporters of neural activity (e.g., GCaMPs) have been successfully used to describe spontaneous sleep in flies (Tainton-Heap et al. 2021; Flores-Valle and Seelig 2022; Bushey, Tononi, and Cirelli 2015), these are typically still limited to a narrow region of interest (e.g., the mushroom bodies, or the central complex), and imaging conditions are rarely commensurate with the typical day-night cycles of normal sleep. In this study, we overcame these drawbacks by recording electrical activity from 16 channels across the fly brain, in behaving flies across long-lasting recordings that spanned a typical day and night. Our multichannel recording preparation therefore approximates as closely as possible - in flies - a sleep EEG, which has been the starting point for most discussions on sleep physiology in other animals. The human sleep EEG has defined the sleep stages that are now being investigated in other animals (Kirszenblat and van Swinderen 2015; Van De Poll and van Swinderen 2021; Raccuglia et al. 2019, 2022), although this is obviously a neocortical view with potentially little relevance to animals lacking the neural architecture giving rise to sleep signatures such as delta (1-4Hz) during slow-wave sleep or theta (5-8Hz) during REM sleep (Jaggard, Wang, and Mourrain 2021).

Rather than focus on specific frequency bands such as delta and theta, we conducted an agnostic analysis of our multichannel LFP data using machine learning techniques. These unbiased classifiers identified distinct stages of sleep, in flies that were otherwise entirely quiescent (apart from certain micro-behaviors, which we discuss further below). These identified sleep stages align closely with similar changes in brain activity dynamics observed in calcium imaging data in spontaneously sleeping flies (Tainton-Heap et al. 2021). For example, in the calcium imaging data we showed that even before sleep onset, the number of ‘active’ neurons is already different (lower) than wake; accordingly, in the current electrophysiological data the classifiers predict sleep onset 2min before flies stop moving. This also aligns with an older (single channel) electrophysiological sleep study in flies showing that brain LFP activity becomes uncorrelated from behavior 5min before sleep onset (B. van Swinderen, Nitz, and Greenspan 2004). Together, these findings make a compelling case for dissociative states in the fly brain, which is consistent with the view that such states might also be changing within a sleep bout.

Our multichannel recordings also revealed that changes in sleep physiology are likely to encompass the entire fly brain, from the optic lobes to the central complex. This is also consistent with other studies, although this has not been previously demonstrated using a comprehensive multichannel approach. An early study in honeybees showed that visually responsive neurons in the optic lobes become unresponsive during sleep (Kaiser and Steiner-Kaiser 1983), and that these cells become rapidly responsive again when bees are woken up with an air puff. Immunochemical studies investigating synaptic proteins found that these were downregulated in the optic lobes during sleep (Donlea, Ramanan, and Shaw 2009), as well as in the whole brain (Gilestro, Tononi, and Cirelli 2009). It is understood that the insect optic lobes receive significant feedback from the central brain, as well as from the contralateral lobes (Scheffer et al. 2020; Mu et al. 2012), and it has been shown that oscillatory neural activity extends throughout the fly brain (Paulk et al. 2013), so our finding that the optic lobes also ‘sleep’ is not quite surprising. Recent work using a similar multichannel recording preparation found that isoflurane anesthesia impacted feedback from the central brain to the optic lobes (Cohen, van Swinderen, and Tsuchiya 2018), suggesting that such efferent communication is a feature of the waking fly brain. Yet, sleep in the central fly brain is different from sleep in the periphery. Interestingly, only central channels were predictive of sleep onset, and only the central channels revealed the 5-10Hz frequency features that we have previously identified in single channel recordings (Yap et al. 2017). This suggests a sleep-regulatory role for the central complex, which aligns well with previous studies (Donlea et al. 2011; Troup et al. 2018; Tainton-Heap et al. 2021).

Sleep in *Drosophila* was originally defined by inactivity criteria, based on locomotion-based readouts (P. J. Shaw et al. 2000; Hendricks et al. 2000). Subsequent studies employing video monitoring and probing arousal thresholds confirmed these simple readouts to be accurate estimates of sleep in flies (van Alphen et al. 2013; Faville et al. 2015; Wiggin et al. 2020), but these behavioral studies also showed that flies slept in distinct stages. Only recently has closer video monitoring of fly micro-behaviors revealed that these animals are not entirely immobile during sleep (van Alphen et al. 2021), although some micro-behaviors were already anecdotally observed in the first reports of fly sleep, such as changes in posture (P. J. Shaw et al. 2000; Hendricks et al. 2000). Other insects, such as honeybees, display characteristic micro-behaviors during sleep, such as changes in posture (Eban-Rothschild and Bloch 2008) and antennal movements (Sauer et al. 2003). Interestingly, in our study we also found evidence of altered antennal movements during fly sleep, alongside the previously reported proboscis extensions (van Alphen et al. 2021). These micro-behaviors are not necessarily correlated, although they do seem to be increased during mid-sleep epochs. PEs have been associated with a deep sleep function (waste clearance) in a previous study (van Alphen et al. 2021), so their occurrence in rhythmic spells during mid-sleep is consistent with that interpretation.

Interestingly, PEs during wake and sleep are electrophysiologically different, even though they are behaviorally identical. We found that the neural signatures of PEs occurring during wake are concentrated in the middle channels, and spread across all frequencies (5-40Hz). It is interesting to note that these middle channels could coincide with the location of neuropils of the antennal mechanosensory and motor center (AMMC). Several studies (Kain and Dahanukar 2015; Kim, Kirkhart, and Scott 2017) have implicated the AMMC as the location of axons of gustatory projection neurons (GPNs) and thus an immediate higher order processing center for taste. Other studies (Flood et al. 2013) have also shown that persistent depolarization of motor command activity of the Fdg (feeding) neurons could also result in PEs. In this context, it is pertinent to note that LFP activity during PE events in the awake periods are higher than those in the ‘awake’ periods without PE events, suggesting a distinct PE signature. But this is not the case for the exact same behaviors during sleep. We found that LFP activity for PEs occurring during sleep bouts are concentrated instead in the central channels and engage primarily the higher frequencies (32-40 Hz). This suggests a distinct control mechanism for PEs occurring during sleep versus wake, with central brain circuits potentially involved in regulating this sleep-related function.

There are obviously several drawbacks to studying sleep physiology in a tethered animal that has been skewered by a recording electrode. Sleep cannot be quite normal in such a preparation. For example, it is possible that the damage caused by the electrode evokes an increased need for repair (Stanhope et al. 2020) and consequently waste clearance (van Alphen et al. 2021), thus increased PE behavior. However, this would also be the case for windows in the brain created for calcium imaging (Tainton-Heap et al. 2021) (and in the latter scenario the proboscis is typically glued in place to prevent brain motion artifacts), so no fly brain recording preparation (yet) can realistically sidestep these concerns. Nevertheless, it is evident that even in this somewhat contrived context, flies do still sleep and their sleep displays unequivocal evidence of distinct stages.

Our study also paves the way for asking fundamental questions about fly sleep in the following fashion. First, the LFP activity of mutant strains (with higher, or lower baseline sleep) could be recorded and its differences across the wild type could be quantified. Second, for understanding and probing the exact spatial patterns of specific sleep stages identified in this study with higher resolution, 2-photon imaging at the whole brain level could be recorded for longer duration (controlled by closed loop detection of events), while optimizing for signal loss with photo bleaching. Third, closed loop techniques could be employed to disrupt sleep either at the PE stage or at other relevant stages to identify behavioral phenotypes, thereby providing casual evidence for function of the specific stage.

Our multichannel data add to the growing realization that the entire insect brain engages in dynamical patterns of activity during both sleep and wake (Tainton-Heap et al. 2021; Troup, Tainton-Heap, and van Swinderen 2023), and does not simply shut off when insects become immobile or quiescent. To understand these patterns of activity and how they might relate to conserved sleep functions (Van De Poll and van Swinderen 2021) requires novel approaches derived from machine learning, as done in this study, rather than approximations inspired from human EEG.

## Materials and Methods

### Animals

Flies (*Drosophila melanogaster*) were reared on a standard fly medium under a 12h light/dark cycle (lights on at 8 A.M). Flies were raised on a 25°C incubator (Tritech research inc) with 50-60% humidity and fewer than 5 flies were maintained per vial to ensure optimal nutrition and growth. Adult female flies (<3 days post eclosion) of wild-type Canton-S (CS) were used for the electrophysiological recordings. The choice of age of flies was based on pilot data that suggested a higher survival rate of younger flies over a 12h period on the air supported ball setup (after electrode insertion). Flies used for the behavioral dataset were between 3 - 7 days post eclosion. For thermogenetic experiments refer to (Yap et al. 2017) for further details.

### Fly tethering

First, flies were anesthetized on a thermoelectric cooled-block maintained at a temperature of 1-2°C. Second, the thorax, dorsal surface and wings of the fly were glued to a tungsten rod using dental cement (Coltene Whaledent Synergy D6 Flow A3.5/ B3) and cured using high intensity blue light (Radii Plus, Henry Scheinn Dental) for about 30-40 sec. Further, dental cement was also applied to the necks to stabilize them and prevent lateral movement of the head during electrode insertion (next section). Third, to prepare the fly for the multichannel overnight recording, we placed a sharpened fine wire made of platinum into the thorax (0.25 mm; A-M systems). The platinum rod serves as a reference electrode and helps filter the noise originating from non-brain sources. The insertion of a platinum electrode (while providing minimal discomfort to movement of animal) was done using a custom holder with a micro-manipulator to enable targeted depth of insertion. For flies in the behavioral dataset, the procedure was the same, except that no reference wire was inserted.

### Multichannel preparation

First, the tethered fly from the previous step was placed on an air supported ball (polystyrene) that served as a platform for walking/rest. Humidified air was delivered to the fly using a tube below the ball (also from the side) to prevent desiccation. Second, to record from half of the regions in the fly brain (half-brain probe) we used a 16-electrode linear silicon probe (model no. A1×16-3 mm50-177; NeuroNexus Technologies). Third, the probe was inserted into the eye of the fly laterally using a micro-manipulator (Merzhauser, Wetzlar, Germany). The probe was inserted such that the electrode sites faced the posterior side of the brain. The final electrode position (depth of insertion) was determined using the polarity reversal procedure described below. For flies recorded in the behavioral dataset the setup was similar, except that a custom chamber was lowered over the ball and fly to maintain a humidified environment during recordings.

### Polarity reversal

Variability in spatial location of recording sites across different flies is a primary impediment when comparing data across different flies. This occurs mainly due to the angle and depth of insertion of the probe, both of which cannot be precisely controlled. To overcome this issue and to obtain comparable recording sites across flies, we designed a novel paradigm using visual evoked potentials (*Figure S2*). First, while the probe was being inserted from the periphery to the center of the brain, we used visual stimuli (square wave of 3 sec in duration with 1Hz frequency) from a blue LED. When the visual stimuli was displayed we simultaneously recorded the local field potentials from the 16 electrode sites. During the initial stage of insertion, most of the electrodes are outside of the brain and only a few are inside the eye, optic lobe. The recordings in the electrodes inside the eye, brain show a visual evoked potential corresponding to the leading edge and the trailing edge of the square wave. Second, we move the probe slowly towards the center of the brain so more of the electrode sites would now be inside the brain. Third, we notice that some electrodes have a negative deflection and some have a positive deflection with respect to the leading edge of the square wave. The electrodes in the eye, optic lobe regions display a positive deflection and electrodes further to the central parts of the brain display a negative deflection. However this polarity change usually happens in the electrodes that are coincident on the regions right after the medulla. Fourth, for all flies we made sure that the polarity change coincided with the electrodes 11-13 inorder to establish consistency in terms of the spatial locations.

### Dye based localization

Inorder to identify the possible locations in the brain targeted by the electrodes, we used a three step procedure. In the first stage, we used immunohistochemistry to identify the locations of electrodes using a fluorescent dye and neuropils using antibodies against nc82 (presynaptic marker bruchpilot) respectively. In the second stage, we used a registration procedure to map the dye locations to an EM dataset (using nc82 images). In the third stage, we used principal component analysis to identify the precise neuropils targeted.

#### a ) Immunohistochemistry

First, we labeled the probe with Texas red fluorescent dye conjugated to 10,000-Da mol mass dextran dissolved in distilled water (Invitrogen) to identify the recording locations. Second, after removing the flies from the tether, the brains were dissected in ice cold 1x phosphate buffer solution (PBS) and fixed in 4% paraformaldehyde diluted in PBS-T (1× PBS, 0.2 Triton-X 100) for 20 minutes in dark to preserve the fluorescence of the dye. Third, after fixation, tissues were washed 3 times with PBS-T (0.2% Triton X-100 in PBS (PBST) with 0.01% sodium azide (Sigma Aldrich)) and blocked for 1 hour in 10 % Goat Serum (Sigma Aldrich). Fourth, the brains were then incubated overnight in a primary antibody solution (mouse anti-nc82 1:20 DSHB). Fifth, on the next day brains were washed 3 times with PBS-T (10 min per wash) and incubated overnight in a secondary anti-body solution (1:250 goat anti-mouse Alexa 647). Finally, the brain was washed in PBST and embedded in Vectashield and imaged using a confocal microscope (Zeiss).

#### b ) Image registration

First, for each fly we used the nc82 image as source space to align to the JFRC2 template space (which is a spatially calibrated version of JFRC (Jenett et al. 2012) from FlyLight). The registration process involved two steps: i) rigid affine registration that roughly aligned the source image to the template space with 12 degrees of freedom (translation, rotation, scaling). ii) non-rigid registration that allowed different brain regions to move independently with a smoothness penalty. The entire process was carried out using the CMTK plugin (FiJi toolbox) as described here (Ostrovsky, Cachero, and Jefferis 2013). Second, we then used the JFRC2 (light-level) registration as bridging registration to FAFB14 (EM dataset) using the natverse toolbox (Bates et al. 2020) and mapped both the nc82 images and the dye locations to the FAFB14 space.

#### c ) Electrode localisation

The electrode dye locations inside the brain are usually visible as fragments (points) instead of a single continuous (line) segment, mainly because the insertion of the probe causes the smearing of the dye on the neuropils in the brain. Inorder to identify the precise locations of the recording electrodes in the brain, we first used the points and performed principal component analysis to find the eigenvector or line (1st principal component) that would have minimize the distance between the different points to the line itself. This line could be thought of as the main path of the probe as it entered into the brain. Next, we choose the innermost electrode as the projection of the innermost point (dye location) projected onto the eigenvector. The rest of the recording electrode sites were obtained by sampling the same eigenvector at intervals of 25 µm (which is the interelectrode distance on the probe) from the innermost point.

### LFP recording

The LFP data from the 16-electrode probe was acquired using Tucker–Davis Technologies (Tucker-Davis Technologies, US) multichannel data acquisition system at 25 kHz coupled with a RZ5 Bioamp processor and RP2.1 enhanced real-time processor. Data was acquired and amplified using a pre-amplifier (RA16PA/RA4PA Medusa PreAmp). The pre-amplifier used can only record data of up to 20 hours on a single charge cycle, hence we limited the recording of the LFP signals to 20 hour duration. Further, as file sizes tend to be larger over longer recording periods, we recorded data in chunks of 1 hour which was automatically controlled via a MATLAB script.

### Video recording for flies on electrophysiology setup

The ball setup was illuminated with visible light, switched ON at 8 AM and switched OFF at 8 PM (mimicking the light/dark cycle conditions in the incubator). Further, we used Infrared LEDs for monitoring the movement of the fly on the ball (which allowed us to quantify movements under both the light and the dark cycle. We recorded the fly in profile view with a digital camera from Scopetek (DCM 130E) and to achieve optical magnification, we used a zoom lens (from Navitar). As done previously (Yap et al. 2017), we removed the IR filter in front of the camera sensor, to allow for filming under IR light, thereby achieving constant illumination under both day and night. We made a custom script with Python (2.7.15), OpenCV (3.4.2.17), that allowed for recording videos automatically and saving them in hourly intervals. The video was recorded with a resolution of 640 x 480 pixels at 30 frames per second using Xvid codec and further with additional metadata (time stamps in a csv file) that allowed a later matching up of the LFP data with the video data.

### Video recording for flies on behavioral dataset setup

The camera in this setup was a Point Grey/Teledyne FLIR Firefly perpendicular to the fly, in addition to an extra camera (ProMicroScan) placed on the trinocular output of a Nikon SZ7 stereomicroscope. This second camera was used to record a close-up view of the head of the fly for the purposes of tracking movements of the antennae. Illumination was as above with infrared LEDs and recordings were obtained with the same Python scripts.

### Movement analysis

The fly movement was quantified with the video files using Python (3.6.1), OpenCV (3.4.9) in the following manner. First, every video file (1 per hour of recording) was read frame by frame. Second, for each frame, we clipped the image such that the main focus was on the fly while ignoring items in the background. Third, we converted the color space for each frame from BGR to grayscale. Fourth, we computed the ‘deltaframe’ as the absolute difference of the current frame with the previous frame. Fifth, we thresholded the deltaframe using a custom defined threshold per fly and converted them into binary. Sixth, we dilated the thresholded image and identified contours in the dilated image and looped over the different contours selecting those above a specific threshold (area). Finally, we drew rectangles around the contours above the threshold on the original (color) image to manually verify the movement location. Only those frames that had contours above threshold were regarded as ‘moved’ frames, other frames would be classified as ‘still’. Thus, each frame would be either 0 (still) or 1 (moved). In the next stage, we used the frame by frame movement data to identify segments of LFP data as ‘sleep’ or ‘awake’ etc in the following fashion. First, we synced the LFP data with the video data by using the time stamps in both the LFP data and video metadata (csv files). Second, we clipped both the LFP and video data to the first 8 hours of recording. Though 23 flies survived for more than 24 hours, we only used the first 8 hours to ensure that the fly’s health was completely optimal (considering the circumstances) in both the behavior and brain recordings. Further only 16 flies were used for the analysis, as 7 of them had issues with calibration (noisy or no calibration) or abnormal activity (either no sleep trials or very active). Third, we pruned movement data to ensure brief noise in movements are avoided. Fourth, we identified the segments of data wherein the fly was immobile for more than 5 mins as ‘sleep’ and the segment immediately preceding 2 mins before the sleep data as ‘presleep’ and the rest of the data as ‘awake’.

### LFP analysis

#### a) Preprocessing

LFP data was analyzed with custom-made scripts in MATLAB (The MathWorks) using EEGLAB toolbox (Delorme and Makeig 2004). The preprocessing steps were as follows: First, the binary data was extracted for every hour from Tucker-Davis technology ‘tank’ file format to MATLAB ‘mat’ file format. Second, the data were resampled to 250 Hz and bandpass filtered with zero phase shift between 0.5 and 40 Hz using hamming windowed-sinc FIR filter, further line noise at 50 Hz was removed using a notch filter. Third, the hourly LFP data was saved to EEGLAB ‘.set’ file format. Fourth, the hourly LFP data were interpolated in a linear way to avoid any discontinuities between the hourly segments of data. Fifth, the movement data (see Movement analysis) was added to the EEGLAB file along with the start and end time based on video data. Sixth, the multi-hour LFP data (along with the movement data) is collated for the first 8 hours of the recording. Seventh, we created separate epochs based on movement data into ‘sleep’, ‘presleep’, ‘awake’ (where preceding 2 mins of immobility (−2 to 0 mins) is ‘presleep’ and immobility is ‘sleep’ and the rest of the data is ‘awake’, here 0 mins is the start of the immobility). Eighth, the epochs were now re-referenced based on the channel where the polarity reversal occurred. For this we identified the channel wherein the polarity reversal occurred (see Polarity reversal section) and subtracted all the channels from this channel, thus resulting in 15 channels after the re-referencing. This brain based referencing technique (similar to the Cz based reference in human EEG recordings) allows for filtering of non-brain based physiological noise (like heartbeat etc). Previous multichannel recordings used only the thorax based referencing (followed by bipolar referencing) along with Independent Component Analysis (ICA) to remove physiological noises. However, the identification of noise components like heartbeat etc from ICA is subjective and further depends on the expertise of the human curator. Our technique overcomes these issues while simultaneously providing a method to remove physiological noises not originating from the brain.

#### b ) Power spectrum analysis (sleep vs awake)

The power spectra of the LFP data was computed for each fly in the following fashion. First, each condition (‘awake’, ‘sleep’ etc) of varying duration was re-epoched into trials of 60 sec duration. Second, each trial was bandpass filtered with zero phase shift between 5 and 40 Hz using hamming windowed-sinc FIR filter. Third, for each trial, power spectra (in decibels) was computed using the ‘spectopo’ function in the EEGLAB toolbox in MATLAB. Fourth, the mean power spectra for all the trials per condition per fly was computed. The goal of the power spectra analysis was to identify the cluster of frequency bands and channels that differ across the sleep, awake periods at the group level. To perform these group level comparisons (sleep vs awake periods) we only used flies that had at least 10 trials under each condition. We performed a cluster permutation test (flies x frequencies x channels) using MNE (0.22.0) in python (permutation_cluster_1samp_test) (Gramfort et al. 2013) with all possible permutations to identify clusters that differ across awake and sleep periods. We also computed the effect sizes for every channel x frequency combination using cohen’s d measure (difference of means/ standard deviation).

### Thermogenetic sleep induction

The thermogenetic sleep induction data was collected using 104y-Gal4 lines as part of the study (Yap et al. 2017). This multichannel recording consisted of a 16-electrode full-brain probe (model no. A1×16-3mm50-177; NeuroNexus Technologies) covering the whole of the brain (*Figure S6B*) (in contrast to the half-brain probe mentioned before) with interelectrode distance of 50 µm. The rest of the recording parameters were the same as mentioned in the previous section. Sleep induction was achieved by transient circuit activation of the sleep promoting circuit innervating the dorsal fan shaped body (dFB). For example, this was done by using the 104y gal4 lines (offering cell type specificity in the dfB regions) to control the expression of UAS driven TrpA1 (temperature sensitive cation channel), thereby allowing for the activation of the specific neurons in dFB with temperature changes. As described in (Yap et al. 2017), before the induction of sleep, the baseline activity was recorded in the ‘baseline’ condition for 3 secs, followed by stimulation in the ‘sleep induction’ condition for 3 secs before returning to recovery for 3 secs.

#### a) Preprocessing

LFP data was analyzed with custom-made scripts in MATLAB (The MathWorks) using EEGLAB as mentioned before. The preprocessing steps were as follows: First, the LFP data per condition (‘baseline’, ‘sleep induction’, ‘recovery’) was converted to EEGLAB ‘.set’ file format with a sampling rate of 1 KHz. Second, the LFP data was re-referenced using a differential approach, wherein nearby channels are subtracted with each other resulting in 15 channels.

#### b) Power spectrum analysis (baseline vs sleep induction)

The power spectra of the LFP data was computed for each fly in the following fashion. First, each condition (‘baseline’, ‘sleep induction’ etc) was reepoched into trials of 1 sec duration. Second, each trial was bandpass filtered with zero phase shift between 5 and 40 Hz using hamming windowed-sinc FIR filter. Third, for each trial, power spectra (in decibels) was computed using the ‘spectopo’ function in the EEGLAB toolbox in MATLAB. Fourth, the mean power spectra for all the trials per condition per fly was computed. The group level comparison was performed using cluster permutation test methods (as described in previous sections) to identify differences in frequency x channels across ‘preheat’ and ‘heaton’ conditions.

### Sleep staging by classifiers

The main goal of this analysis was to use classifiers to identify the existence of sleep stages using LFP data.

#### a) Labeling of sleep states

Here, we relabelled the segments of data (already identified as ‘sleep’, ‘awake’ based on movement data) in the following fashion. First, we labeled the segments of data in the first 2 mins (0 to 2 mins) after the start of immobility as ‘earlysleep’ and the segments of the data in the preceding 2 mins (−2 to 0 mins) as ‘presleep’. Second, we labeled the segments of data in the last 2 mins of sleep as ‘latesleep’ and the segments of data in between the ‘earlysleep’ and ‘latesleep’ as ‘midsleep’. The rest of the data is considered as ‘awake’.

#### b) Preprocessing & power spectrum computation

The preprocessing steps were the same as mentioned in the previous section (LFP preprocessing). For the computation of the power spectrum, we followed similar procedures as mentioned before, however we saved the individual power spectrum per trial (channels x frequency) per fly in a csv file along with the corresponding label of the sleep state.

#### c) Classifier probability analysis

We implemented a support vector machine (svm) based classifier using scikit-learn (0.24.2) to classify the LFP data using the following steps. First, we collated the features based on power spectrum (channels x frequency) from all the flies across different sleep states. Second, we filtered the features to only ‘awake’ (5106 epochs) and ‘midsleep’ (1165 epochs) states. Here, we also did not feed (for training) the preceding 2 mins of ‘presleep’ and succeeding 2 mins of ‘earlysleep’ and the last 2 mins of sleep ‘latesleep’ into the classifier (we used those minutes for sanity check purposes - Refer to Figure 5A). Third, we encoded the target labels (‘awake’, ‘midsleep’) into binary states using ‘LabelEncoder’ from scikit-learn. Fourth, we balanced the composition of labels (or classes) to prevent bias due to unequal distribution of classes in the training dataset. Fifth, we divided the dataset into train and test sets (80% train, 20% test) using ‘train_test_split’ from scikit-learn in a stratified fashion. Sixth, we subjected both the train and test data to a standard scaler using ‘StandardScaler’ from scikit-learn, which removes the mean of the data and scales it by the variance. Seventh, we implemented a svm based classifier using a ‘linear’ kernel along with probability estimates per class and fit the classifier to the train dataset. Eighth, we used the trained classifier on the test dataset and computed different metrics of classifier performance like accuracy, roc_auc, recall, precision, f1-score etc using ‘metrics’ from scikit-learn (*Figure S7B*). Ninth, we used the trained classifier on all class labels (‘awake’, ‘presleep’, ‘earlysleep’, ‘midsleep’, ‘latesleep’, preceding 2 mins of ‘presleep’ and succeeding 2 mins of ‘latesleep’) from the original dataset and computed the probability estimates per class. It is pertinent to note that none of the ‘presleep’, ‘earlysleep’, ‘latesleep’, preceding 2 mins of ‘presleep’ and succeeding 2 mins of ‘latesleep’ the data have not been seen by the classifier beforehand. The above process from Step 5 onwards is repeated a further 4 times with different test, train splits to create five different iterations of classifiers and performance metrics.

#### d) Multiclass svm analysis & Feature importance

To identify differences across multiple classes (‘awake’, ‘presleep’, ‘earlysleep’, ‘midsleep’, ‘latesleep’) we implemented a random forest classifier using scikit-learn (0.24.2) to classify the LFP data using the following steps. First, we collated the features based on power spectrum (channels x frequency) from all the flies across different sleep states. Second, as the different labels (or classes) were unbalanced viz: ‘awake’(5585 epochs), ‘presleep’(258 epochs), ‘earlysleep’(262 epochs), ‘midsleep’ (1165 epochs), ‘latesleep’ (262 epochs), we used SMOTE (Synthetic Minority Over-sampling Technique) from imblearn (0.8.1) to balance the distribution of classes in the dataset. Third, we divided the dataset into train and test sets (80% train, 20% test) using ‘train_test_split’ from scikit-learn in a stratified fashion. Fourth, we subjected both the train and test data to a standard scaler using ‘StandardScaler’ from scikit-learn, as mentioned in the previous section. Fifth, we encoded the target labels into binary states using ‘LabelBinarizer’ from scikit-learn. Sixth, we implemented a random forest classifier for this multiclass classification problem. As the random forest classifier has multiple hyperparameters that need to be tuned, we first used a random grid (using ‘RandomizedSearchCV’ from scikit-learn) to search for the hyperparameters and then further used these parameters in a grid search model (using ‘GridSearchCV’ from scikit-learn) to identify the best hyperparameters. Seventh, we used the trained classifier on the test dataset and computed different metrics of classifier performance like recall, precision, f1-score etc using ‘metrics’ from scikit-learn separately for all the 5 classes. Furthermore, we also computed a normalized confusion matrix using ‘confusion_matrix’ from scikit-learn. The above process from Step 5 onwards is repeated a further 4 times with different test, train splits to create five different iterations of classifiers and performance metrics. Finally to identify and rank the importance of different features we utilized the permutation importance metric (using ‘permutation_importance’ from scikit-learn). The permutation feature importance works by randomly shuffling a single feature value and further identifying the decrease in the model score (Breiman 2001). The process breaks the relationship between the shuffled feature and the target, thus if the feature is very important, it would be indicated by a high drop in model score, on the other hand if it is relatively unimportant, then the model score would not be affected so much. We used the permutation importance with a repeat of 5, and for each train/test split we computed a permutation importance score. Finally, the mean permutation importance score was computed using all the splits.

#### e) Classifier metrics

The performance of the above-mentioned classifiers (both SVM based, random forest based) was evaluated using metrics like accuracy, recall, precision, roc_auc, f1-score. The definition of these metrics are as follows:

*Recall*: This refers to the ability of a classifier to correctly detect the true class of the epoch among the classifications made. It is obtained by the (TP/TP + FN). It is also known as sensitivity. TP: True Positives, FN: False Negatives.

*Precision*: This refers to the exactness of the classifier. It is obtained by the (TP/TP + FP). TP: True Positives, FP: False Positives.

*F1-score*: This refers to the harmonic mean between precision and recall.

*roc_auc*: This refers to the area under the receiver operating curve. In general, it refers to how efficient the classifier is in identifying different epochs. Scores closer to 1 indicate a highly efficient classifier whereas those closer to 0 indicate otherwise.

*Accuracy*: This is defined as the number of correctly classified epochs divided by the overall number of epochs classified.

*Confusion matrix*: This enables visualization of the classifier performance, by tabulating the predicted classes against actual classes. For multiclass problems (random forest classifiers here), the values in the diagonal indicate where the predicted and actual classes converge, whereas those on the off-diagonal indicate misclassifications.

### Proboscis tracking for flies on electrophysiology setup

#### a) Pose detection

We used DeepLabCut (Mathis et al. 2018) to track the different body parts of the fly using an artificial neural network trained in the following fashion. First, we extracted frames from: sample videos wherein the fly performs the following: normal walking movement on the ball (‘all_body’), proboscis extension periods (’proboscis’) both while asleep and awake. For each fly we extracted videos of the above mentioned categories for the purpose of creating annotation labels. Second, we extracted frames from these videos and further labeled the different body parts: eye, proboscis, leg1_tip, leg1_joint, leg3_tip, leg3_joint, abdomen (Figure 7). Third, we trained the neural network per fly using this dataset with ‘resnet_50’ weights until the loss parameter during training stabilizes. The performance of the network per fly (train, test error in pixels) was in general similar in both the train and test datasets. Fourth, we evaluated the annotation performance manually by labeling a test video and verifying the same. Finally, this trained network (per fly) was used for annotating the video for the first 9 hours of the recording.

#### b) Pose analysis

In the next step, we use the pose detection output to design a classifier capable of identifying proboscis extension periods. First, we manually detected several sample time points (to be used as ground truth for training/testing the classifier) in the video of each fly and identified proboscis time periods and saved them in a ‘csv’ file. Second, we used the pose tracking data (x,y,likelihood) for the body parts of the proboscis, leg1_tip, leg1_joint, eye, abdomen and further computed low pass filtered data (0.1 Hz butterworth filter) of each body part. Further we also computed the moving average (window length of 5 samples) of the filtered data. Third, we computed ‘dist_eyeprob’ as the euclidean distance between the proboscis and eye body part and finally multiplied the same with the likelihood of the proboscis body part. Fourth, we used the above-mentioned body parts (and its derivatives) as features and used the ‘StandardScaler’ from scikit-learn for normalizing the data. Fifth, we divided the dataset into train and test sets (70% train, 30% test) using ‘train_test_split’ from scikit-learn. Sixth, we implemented a svm based classifier using a ‘rbf’ kernel and fit the classifier to the train dataset. Seventh, we used the trained classifier on the test dataset and computed different metrics of classifier performance like accuracy, recall, precision etc using ‘metrics’ from scikit-learn. The data segments (frames) identified here will be used to construct the candidate proboscis periods, which then will be further refined in the next steps.

#### c) Proboscis detection

First, we use the frames identified by the classifier from the previous section and construct continuous segments to identify time periods of probable proboscis periods. Further, we add additional time periods by using the likelihood of the proboscis part with a threshold based method. Second, we identify the peak frame (where the maximum displacement of the proboscis occurs) in each proboscis extension event (each proboscis bout consists of multiple proboscis extension events) and save the identified proboscis events (frame number, time, behavior state) to a ‘csv’ file. Third, each event in the csv file is manually verified and only true events are further taken forward. This process is repeated for all the flies and the proboscis detection accuracy per fly is plotted in Figure 7C.

### Micro-behaviour tracking for flies on behavioral dataset setup

Here, the same method for tracking micro-behaviors via DeepLabCut was used, focusing on the proboscis and abdomen for the lateral camera view (See above), and the base and tip of the left and right antennae for the dorsal view of the fly head. The data from these two streams was imported into a custom MATLAB (2020a) script, which performed synchronization based on the integrated timestamps. After preprocessing, antennal tracking with DeepLabCut was converted into an angle for both respective antennae by calculation of the respective positions of the bases and tips, with the angle of the fly’s head with respect to the camera automatically derived from this data and used to correct the angle of the antennae. For the proboscis a median position was calculated for each recording - assumed to be the resting position - and the distance and angle between the proboscis at any given time point and this median position was calculated. Extensions of the proboscis were derived from this distance data with the ‘findpeaks’ function in MATLAB, with a number of exclusion criteria applied to remove tracking artifacts. For example, detected peaks that exceeded a biologically plausible distance threshold, lasted only for a single frame, or had an implausible instantaneous rise time were excluded. Since this method could potentially be biased towards identifying proboscis activity that follows a prototypical shape, we also employed an alternative proboscis event detection based purely on the current distance of the proboscis from resting. In this we used a manually set threshold for each fly to detect portions in the recording when the proboscis was extended versus not, and for these ‘events’ we calculated the duration and median angle of the proboscis during the span of the event. Periods of antennal periodicity in recordings were calculated based on a Fast Fourier Transform (FFT), applied to time segments of recordings. Since proboscis activity was not sinusoidal in nature (and thus would behave poorly if subjected to an FFT), periodicity for this organ was calculated manually as a factor of timing between individual PEs in that proboscis extensions were periodic if they occurred less than 6s after a preceding proboscis extension. This value was selected from observation of typical inter-PE intervals.

### LFP analysis - proboscis

The main goal of this analysis was to identify the spectral signatures associated with the proboscis extension periods across ‘awake’ and ‘sleep’ states in the LFP data.

#### a) Identification of proboscis periods

First, we used the csv file containing frame by frame detection of manually verified proboscis events (from the section above). Second, we identify periods of proboscis extensions which are close together (within 10 sec of each other) and label them as continuous periods. Third, we add activity labels like ‘awake’ (awake periods without any proboscis activity), ‘awakeprob’ (awake periods with proboscis activity), ‘sleep’ (sleep periods without any proboscis activity), ‘sleepprob’ (sleep periods with proboscis activity), ‘presleep’ (presleep periods without any proboscis activity), ‘presleepprob’ (presleep periods with proboscis activity) based on annotated behaviors. Fourth, we extract the LFP data corresponding to the different time periods across each fly.

#### b) Power spectrum analysis

The preprocessing steps for the extracted LFP data were the same as mentioned in the previous section (LFP preprocessing). For the computation of the power spectrum, we followed similar procedures as mentioned before, however we computed the individual power spectrum per trial (channels x frequency) per fly by re-epoching them into trials of 1 sec in duration (instead of the 60 sec periods for sleep analysis, as the proboscis periods are usually shorter). Then the mean power spectrum for all the trials per condition per fly was computed. Next, we performed cluster permutation tests (flies x frequencies x channels) for identifying the differences across frequencies and channels across different conditions. For this analysis we only used flies that had at least 50 trials under each condition.

### Multilevel models

#### a) Models for antennal, proboscis periodicity

We defined 2 different multilevel models (*Supplementary Table 1,3,5 - left, right antenna, proboscis*) to understand how the likelihood of periodicity varies by sleep epoch. In the null model, the periodicity depends only on the mean per fly (fixed effect) and the fly ID (random effect). In the second model (epoch model), the periodicity depends only on the epoch (fixed effect) and the fly ID (random effect).

These models were fit using the ‘lmer’ function (‘lmerTest’ package) in R (Kuznetsova, Brockhoff, and Christensen 2017) and the winning model is identified as the one with the highest log-likelihood by comparing it with the null model, and performing a likelihood ratio chi-square test (χ2). Finally the winning model was analyzed using the ‘anova’ function (*Supplementary Table 2,4,6 - left, right antenna, proboscis*) in R (Fox and Weisberg 2018).

#### b) Models for spectral analysis

We defined 4 different multilevel models (*Supplementary Table 7*) to understand the modulation of the power spectrum by sleep epoch and channel type. In the null model, the power spectrum depends only on the mean per fly (fixed effect) and the fly ID (random effect). In the second model (epoch model), the power spectrum depends only on the LFP epoch type (fixed effect) and the fly ID (random effect). In the third model (channel model), the power spectrum depends only on the channel type (fixed effect) and the fly ID (random effect). In the fourth model (epoch-channel model), the power spectrum depends on a combination of the LFP epoch type and the channel type, both used as fixed effects, and the fly ID (random effect). These four models were fit using the ‘lmer’ function (‘lmerTest’ package) in R (Kuznetsova, Brockhoff, and Christensen 2017) and the winning model is identified as the one with the highest log-likelihood by comparing it with the null model, and performing a likelihood ratio chi-square test (χ2). Finally the top two winning models were compared against each other using ‘anova’ function in R (Fox and Weisberg 2018), to validate whether the winning model (if it is more complex) is actually better than the losing model (if it is simpler). The epoch-channel model emerged as the winning model, indicating an important contribution from different channels. The epoch-channel was further analyzed with the ‘anova’ function (*Supplementary Table 8*) in R (Fox and Weisberg 2018)

#### c) Models for PE event counts

We defined 2 different multilevel models (*Supplementary Table 9*) to understand the modulation of PE event count by sleep epochs. In the null model, the PE event count depends only on the mean per fly (fixed effect) and the fly ID (random effect). In the second model (time_label model), the PE event count depends only on the specific temporal sleep stage (fixed effect) and the fly ID (random effect). These 2 models were fit using the ‘lmer’ function (‘lmerTest’ package) in R (Kuznetsova, Brockhoff, and Christensen 2017) and the winning model is identified as the one with the highest log-likelihood by comparing it with the null model, and performing a likelihood ratio chi-square test (χ2). Thus, the time_label model emerged as the winning model. The time_label model was further analyzed with the ‘anova’ function (*Supplementary Table 10*) in R (Fox and Weisberg 2018)

## AUTHOR CONTRIBUTIONS

Conceptualization: S.R.J., R.J., B.v.S.; Data Curation: S.R.J., R.J., and M.V.D.P.; Formal Analysis: S.R.J., and M.V.D.P.; Investigation: R.J., and M.V.D.P.; Visualization: S.R.J., and M.V.D.P.; Software: S.R.J.; Writing – original draft: S.R.J., and B.v.S.; Writing – review & editing: S.R.J., R.J., M.V.D.P., B.v.S.; Methodology: S.R.J., R.J., M.V.D.P., B.v.S.; Resources: R.J., M.V.D.P., B.v.S.; Project Administration: S.R.J., R.J., B.v.S.; Supervision: B.v.S.; Funding acquisition: S.R.J., and B.v.S.;

## Acknowledgements

The authors would like to thank Dr. Deniz Ertekin for performing immunostaining to identify electrode locations. This work was supported by the Gates Cambridge Scholarship, fieldwork funding from the Department of Psychology to S.R.J.; NIH RO1 NS076980-01 to BvS and Paul Shaw (Washington University); NHMRC GNT1164499 to BvS. We thank members of the van Swinderen lab for critical discussions and Dr. Melvyn Yap for initial code/dataset sharing. S.R.J is grateful to Dr. Tristan Bekinschtein for his continuous encouragement and support which he received while shifting to the field of fly neurobiology.

## Supplementary material

**Suppl Table 1:** Model comparison - Left antenna

| Model | Parameters | Log-likelihood | Pr(>$\chi^2$) |
| --- | --- | --- | --- |
| Null | Fixed: mean, Random: fly ID | -106.53 | - |
| epoch | Fixed: time_label, Random: fly ID | **-73.58** | <0.001 |

**Suppl Table 2:** Type III analysis of variance with Satterthwaite’s method of the winning model (Epoch) - Left antenna

| Model elements | Sum Sq | Mean Sq | NumDF | DenDF | F value | Pr(>F) |
| --- | --- | --- | --- | --- | --- | --- |
| Epoch | 4.5527 | 1.1382 | 4 | 1090 | 16.985 | <0.001 |

**Suppl Table 3:** Model comparison - Right antenna

| Model | Parameters | Log-likelihood | Pr(>$\chi^2$) |
| --- | --- | --- | --- |
| Null | Fixed: mean, Random: fly ID | -68.415 | - |
| epoch | Fixed: time_label, Random: fly ID | **-44.468** | <0.001 |

**Suppl Table 4:** Type III analysis of variance with Satterthwaite’s method of the winning model (Epoch) - Right antenna

| Model elements | Sum Sq | Mean Sq | NumDF | DenDF | F value | Pr(>F) |
| --- | --- | --- | --- | --- | --- | --- |
| Epoch | 3.1004 | 0.7751 | 4 | 1125 | 12.232 | <0.001 |

**Suppl Table 5:** Model comparison - PEs

| Model | Parameters | Log-likelihood | Pr(>$\chi^2$) |
| --- | --- | --- | --- |
| Null | Fixed: mean, Random: fly ID | -301.09 | - |
| epoch | Fixed: time_label, Random: fly ID | **-207.25** | <0.001 |

**Suppl Table 6:** Type III analysis of variance with Satterthwaite’s method of the winning model (Epoch) - PEs

| Model elements | Sum Sq | Mean Sq | NumDF | DenDF | F value | Pr(>F) |
| --- | --- | --- | --- | --- | --- | --- |
| Epoch | 20.877 | 5.2192 | 4 | 795 | 52.923 | <0.001 |

**Suppl Table 7:** Model comparison - LFP power spectrum

| Model | Parameters | Log-likelihood | Pr(>$\chi^2$) |
| --- | --- | --- | --- |
| Null | Fixed: mean, Random: fly ID | -68117 | - |
| Epoch | Fixed: epoch, Random: fly ID | -67941 | <0.001 |
| Channel | Fixed: channel, Random: fly ID | -52391 | <0.001 |
| Epoch-Channel | Fixed: epoch*channel, Random: fly ID | **-51593** | <0.001 |

**Suppl Table 8:**
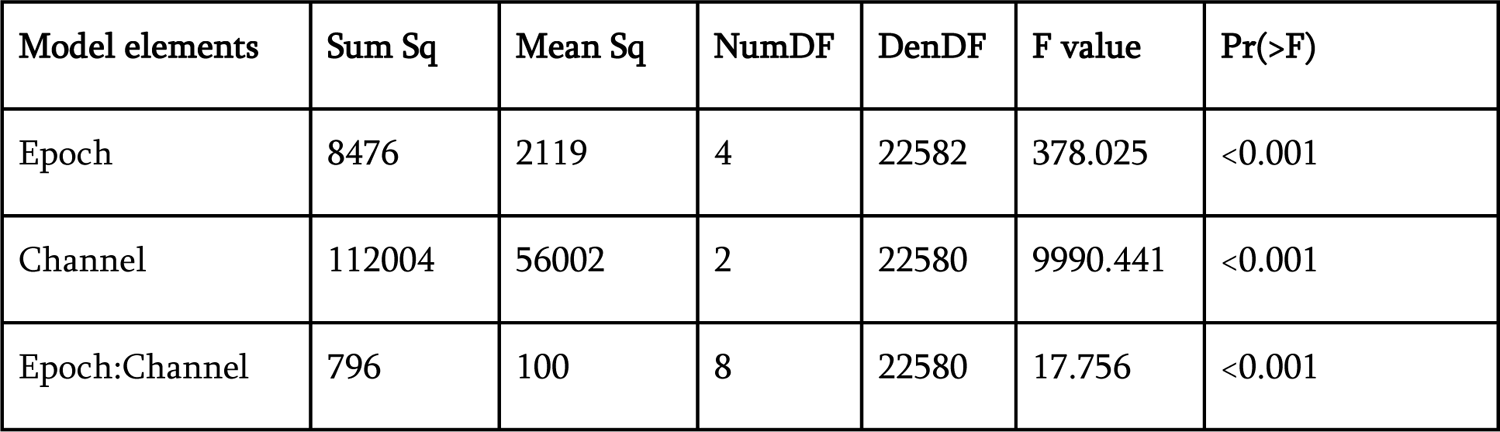
Type III analysis of variance with Satterthwaite’s method of the winning model (Epoch- Channel) - LFP power spectrum

| Model elements | Sum Sq | Mean Sq | NumDF | DenDF | F value | Pr(>F) |
| --- | --- | --- | --- | --- | --- | --- |
| Epoch | 8476 | 2119 | 4 | 22582 | 378.025 | <0.001 |
| Channel | 112004 | 56002 | 2 | 22580 | 9990.441 | <0.001 |
| Epoch:Channel | 796 | 100 | 8 | 22580 | 17.756 | <0.001 |

**Suppl Table 9:** Model comparison - PEs LFP dataset

| Model | Parameters | Log-likelihood | Pr(>$\chi^2$) |
| --- | --- | --- | --- |
| Null | Fixed: mean, Random: fly ID | -4.5588 | - |
| time_label | Fixed: time_label, Random: fly ID | **15.0632** | <0.001 |

**Suppl Table 10:** Type III analysis of variance with Satterthwaite’s method of the winning model (time_label) - PEs LFP dataset

| Model elements | Sum Sq | Mean Sq | NumDF | DenDF | F value | Pr(>F) |
| --- | --- | --- | --- | --- | --- | --- |
| time_label | 1.2145 | 0.20241 | 6 | 41 | 9.6039 | <0.001 |

**Supplementary Figure 1:**
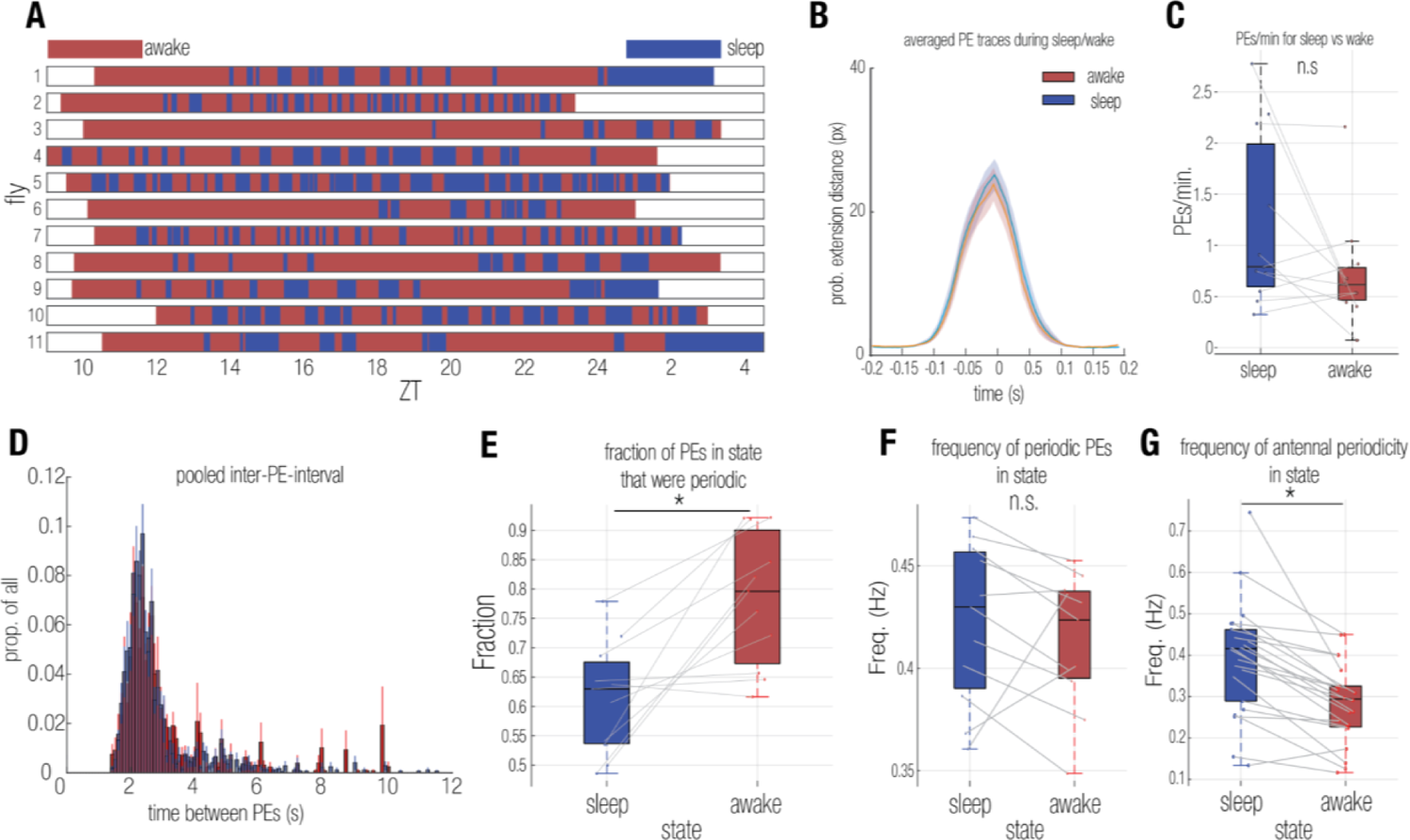
Additional metrics of proboscis activity during sleep and wake. A) Representation of the distribution of sleep (Blue) and wake (Red) across N=11 recorded individuals over the course of time. B) Averaged timecourse of proboscis extension distance from resting during a single event for sleep (Blue) and wake (Red). C) Comparison of proboscis extension rates during sleep and wake (n.s.; Student’s T-test). D) Histogram of the distribution of times between PEs during sleep (Blue) and wake (Red). E) Comparison of the fraction of PEs that were periodic versus isolated for sleep and wake (*p* < 0.05; Student’s T-test). F) Comparison of the average frequency of PE periodicity across sleep and wake (n.s.; Student’s T-test). G) As with F, for antennal periodicity (*p* < 0.05; Student’s T-test).

**Supplementary Figure 2:**
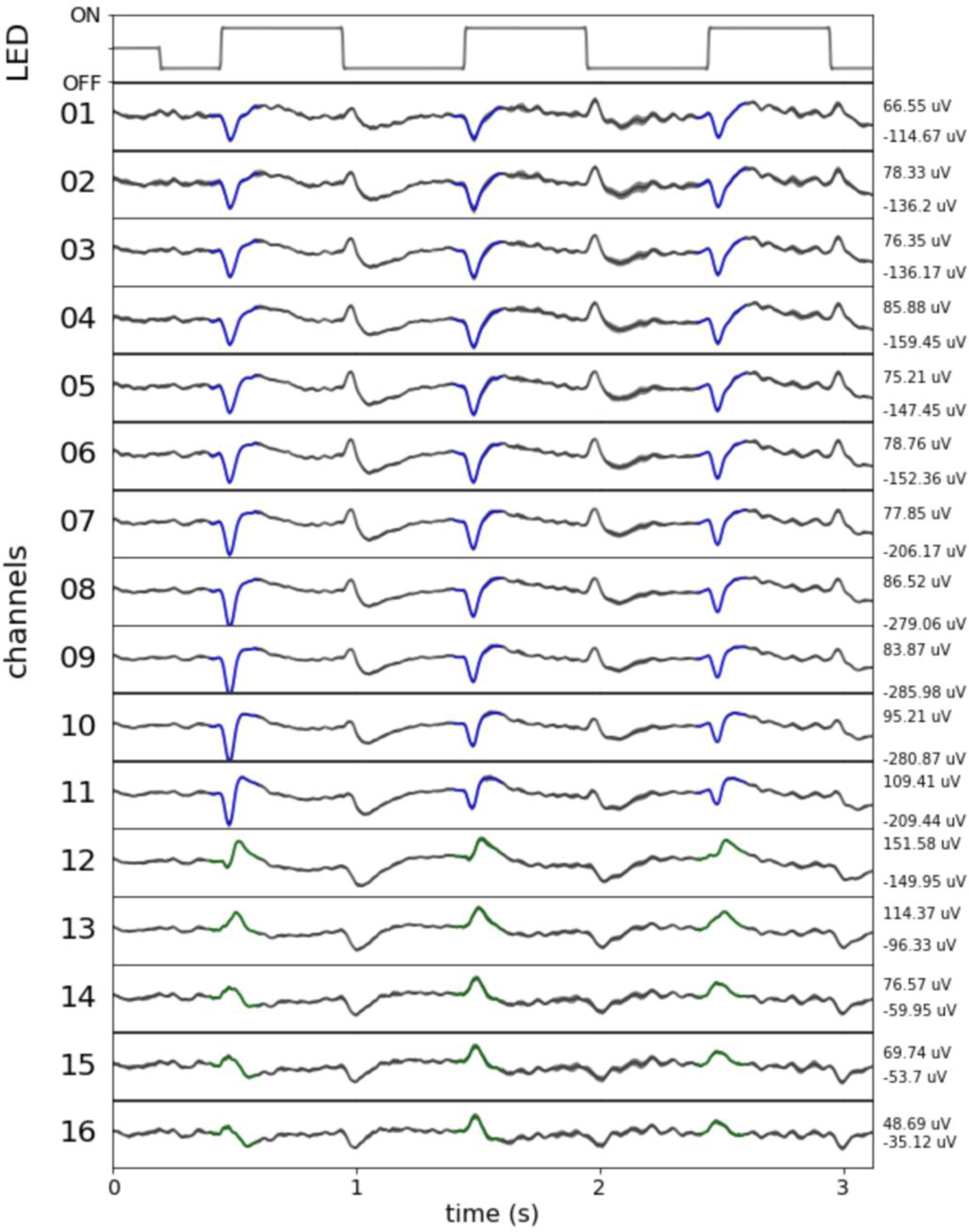
Electrode insertion depth was controlled by using a polarity reversal method. In this example fly, the change in LED stimulation (OFF to ON) stage, coincides with a LFP deflection. The LFP deflection changes from positive (12th channel) to negative (11th channel). The LFP amplitude depicted here is based on an average of 5 trials, with the shaded region representing the standard error.

**Supplementary Figure 3:**
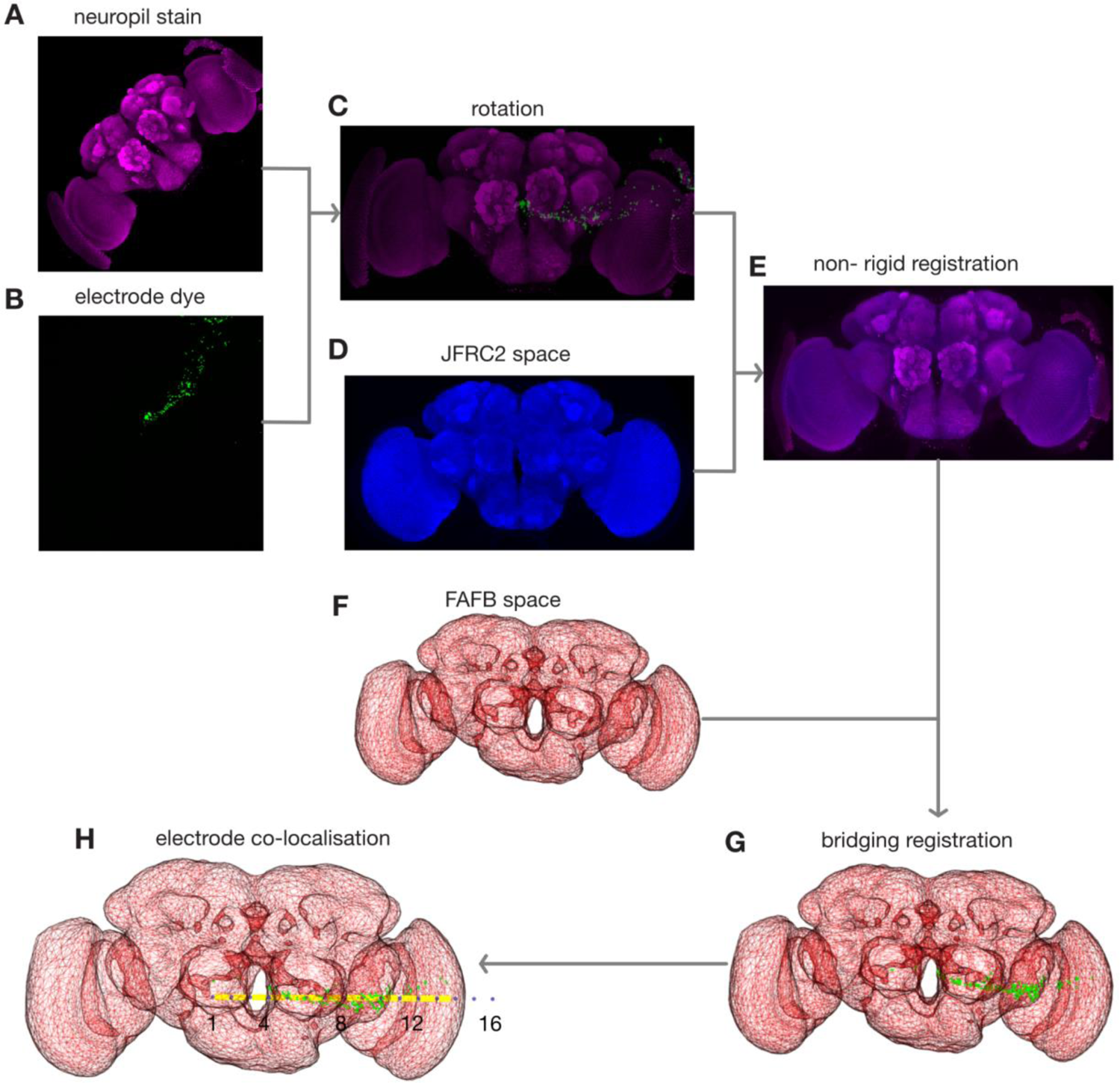
Electrode locations were determined using a dye based localisation method. Neuropil stain (A) and electrode dye locations (B) were registered to JFRC2 space (D) via non-rigid registration. Further Bridging registration was used to register to FAFB space (F), the registration templates were applied on electrode dye locations (B) to produce co-localisation (H).

**Supplementary Figure 4:**
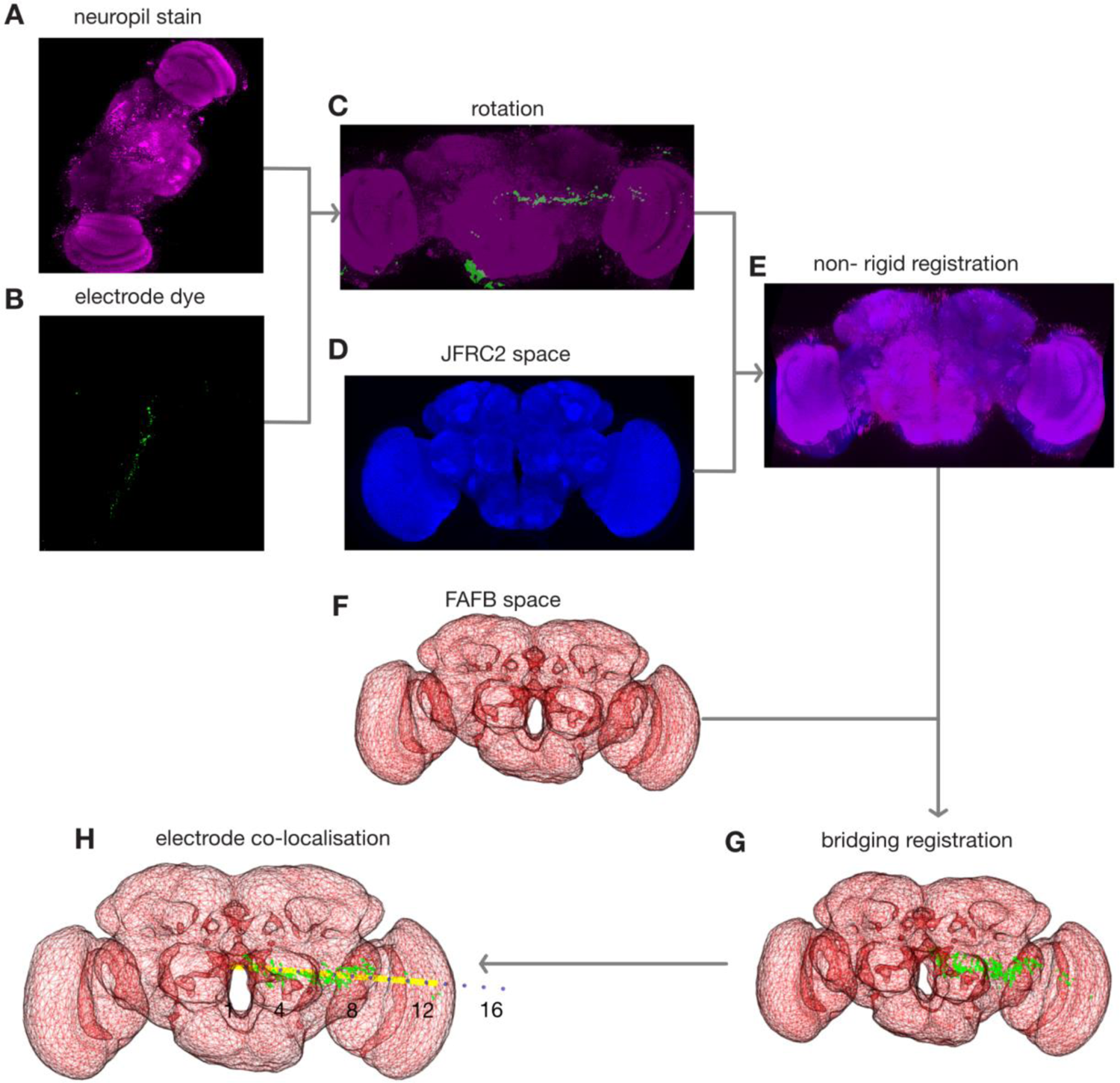
Electrode locations were determined using a dye based localisation method. Neuropil stain (A) and electrode dye locations (B) were registered to JFRC2 space (D) via non-rigid registration. Further Bridging registration was used to register to FAFB space (F), the registration templates were applied on electrode dye locations (B) to produce co-localisation (H).

**Supplementary Figure 5:**
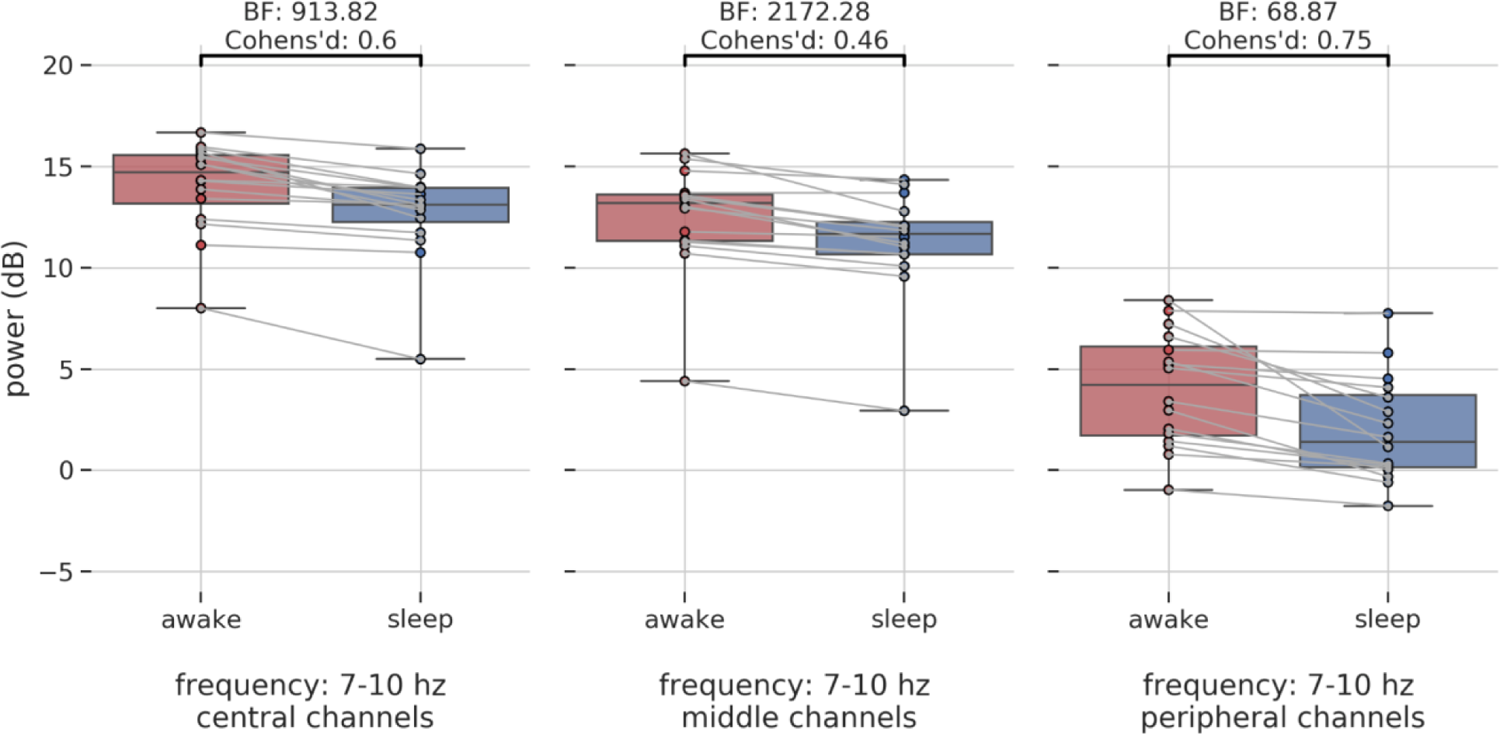
Power differences across central, middle, peripheral channels in the frequency bands of 7-10 Hz.

**Supplementary Figure 6:**
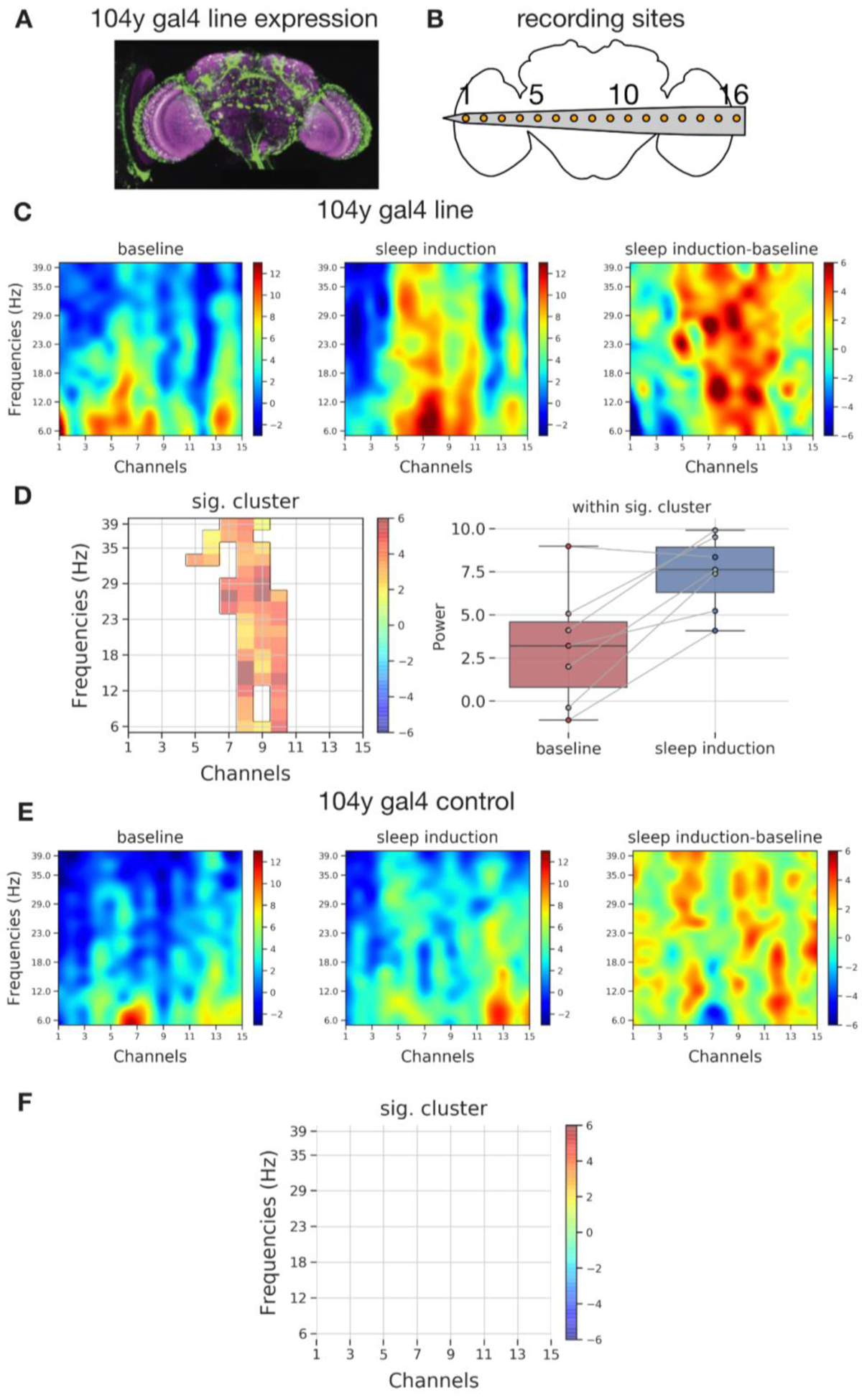
Spectral differences in thermogenetically induced sleep recorded using full brain probe.

**Supplementary Figure 7:**
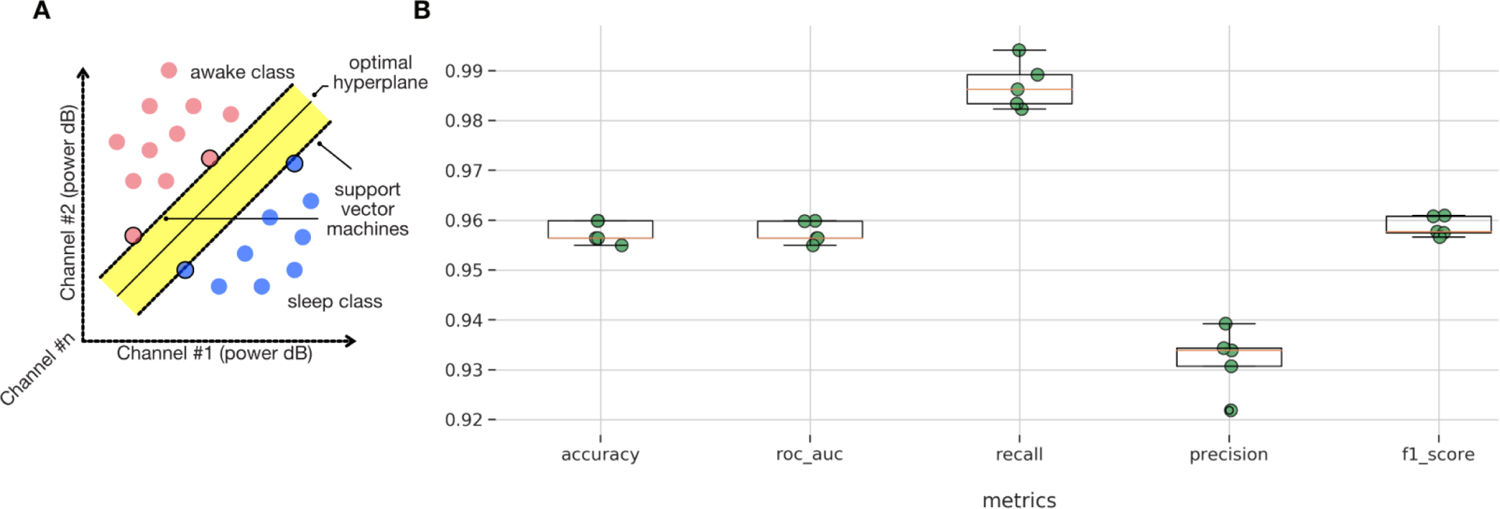
A) Schematic indicating the optimal separation of awake and sleep classes using classifiers based on support vector machines. B) SVM based classifier performance across different metrics based on 5 different train/test data splits.

**Supplementary Figure 8:**
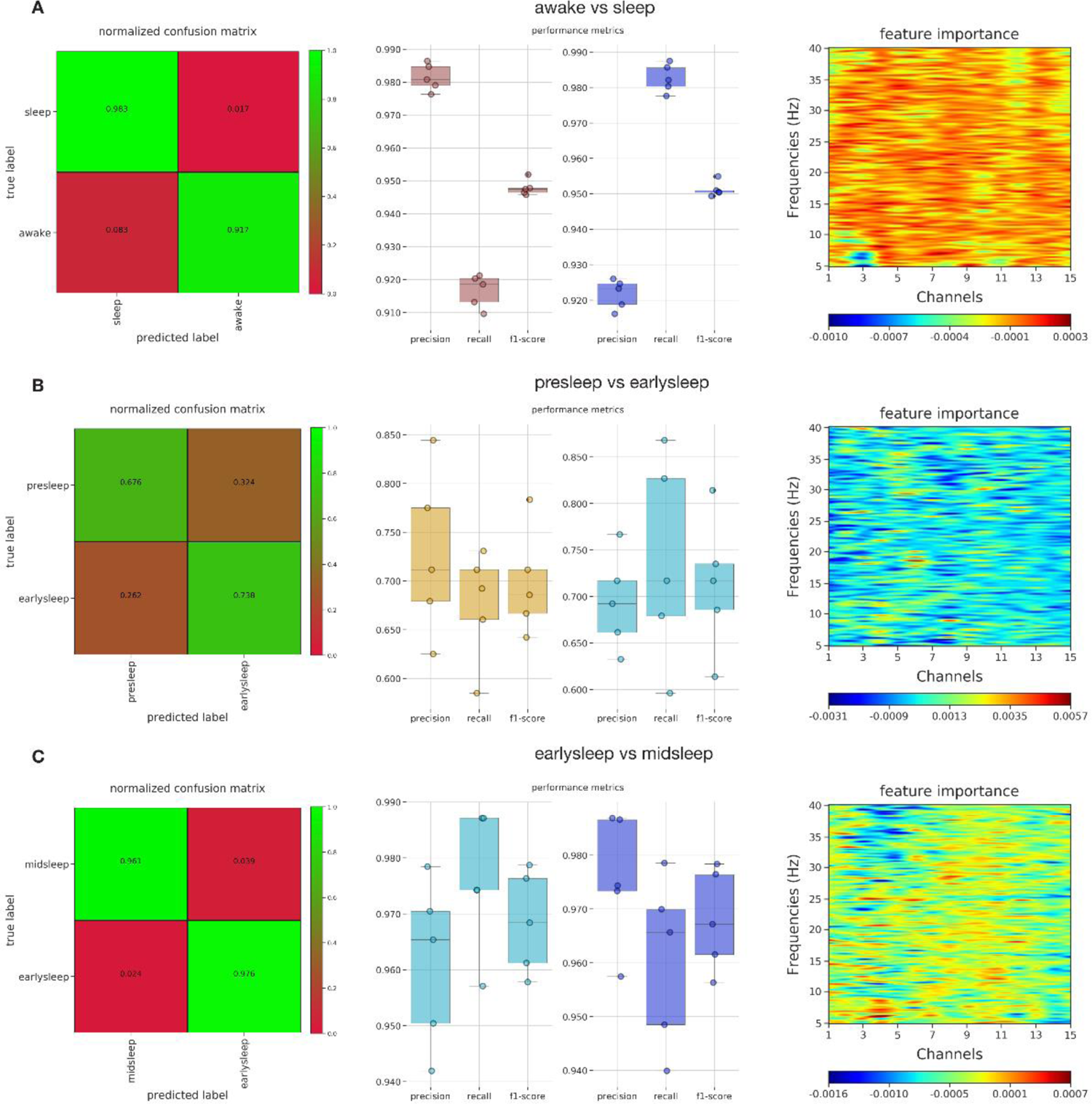
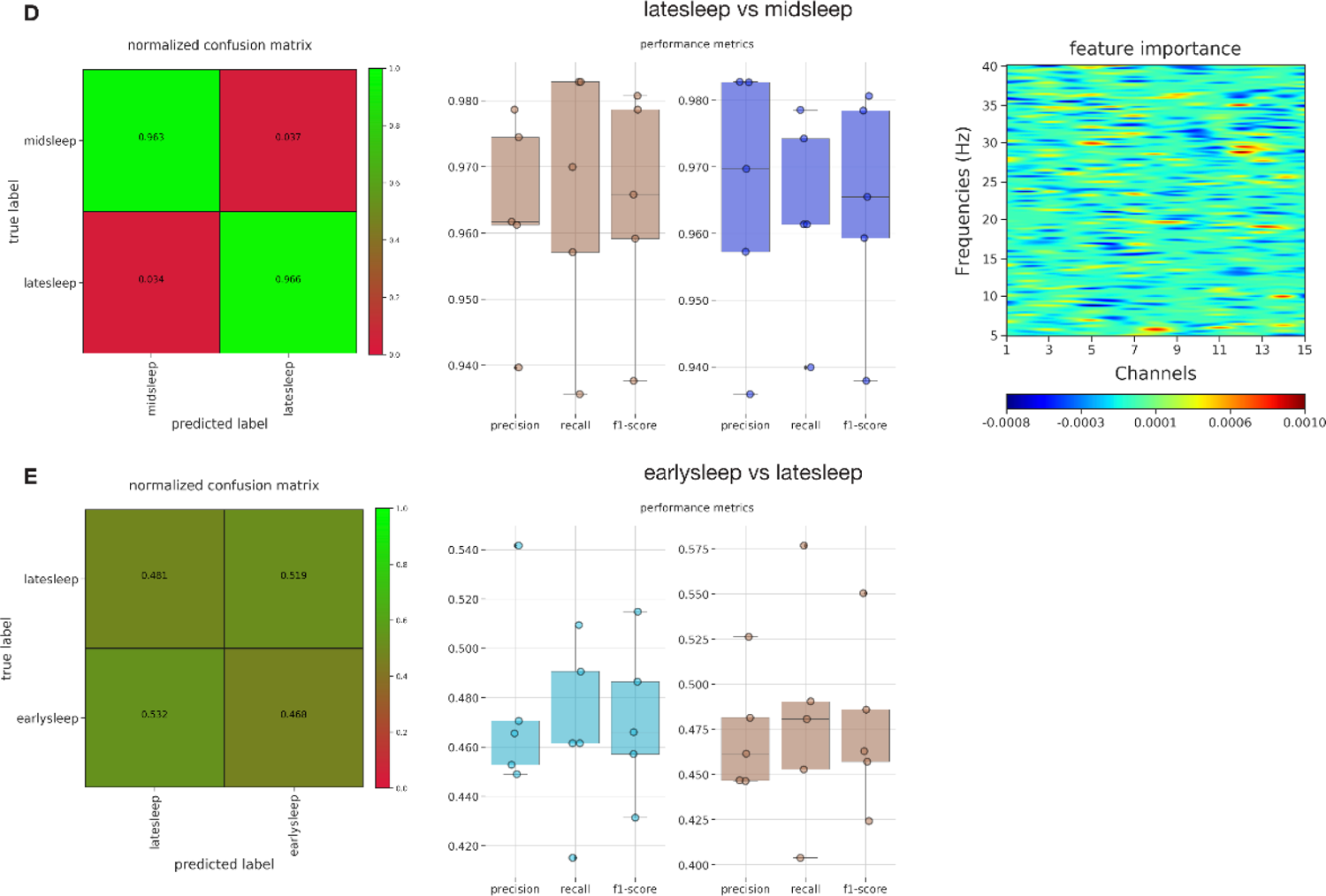
A) Feature importance of the multiclass classifier (reduced to awake vs sleep) indicates an ROI across all channels and almost all frequency bands as critically important. This cross validates the differences in the power spectrum across awake and sleep as shown in Figure 4D. B,C,D,E) Feature importance of multiclass classifier for the other categories.

## References

van Alphen, Bart, Evan R. Semenza, Melvyn Yap, Bruno van Swinderen, and Ravi Allada. 2021. “A Deep Sleep Stage in *Drosophila* with a Functional Role in Waste Clearance.” Science Advances. https://doi.org/10.1126/sciadv.abc2999.

van Alphen, Bart, Melvyn H. W. Yap, Leonie Kirszenblat, Benjamin Kottler, and Bruno van Swinderen. 2013. “A Dynamic Deep Sleep Stage in Drosophila.” The Journal of Neuroscience: The Official Journal of the Society for Neuroscience 33 (16): 6917–27.

Anthoney, Niki, Lucy A. L. Tainton-Heap, Hang Luong, Eleni Notaras, Qiongyi Zhao, Trent Perry, Philip Batterham, Paul J. Shaw, and Bruno van Swinderen. 2023. “Experimentally Induced Active and Quiet Sleep Engage Non-Overlapping Transcriptomes in.” *bioRxiv: The Preprint Server for Biology*, April. https://doi.org/10.1101/2023.04.03.535331.

Bates, Alexander Shakeel, James D. Manton, Sridhar R. Jagannathan, Marta Costa, Philipp Schlegel, Torsten Rohlfing, and Gregory Sxe Jefferis. 2020. “The Natverse, a Versatile Toolbox for Combining and Analysing Neuroanatomical Data.” eLife 9 (April). https://doi.org/10.7554/eLife.53350.

Besedovsky, Luciana, Tanja Lange, and Jan Born. 2012. “Sleep and Immune Function.” Pflügers Archiv - European Journal of Physiology. https://doi.org/10.1007/s00424-011-1044-0.

Breiman, Leo. 2001. “Random Forests.” Machine Learning 45 (1): 5–32.

Bushey, Daniel, Giulio Tononi, and Chiara Cirelli. 2015. “Sleep- and Wake-Dependent Changes in Neuronal Activity and Reactivity Demonstrated in Fly Neurons Using in Vivo Calcium Imaging.” Proceedings of the National Academy of Sciences of the United States of America 112 (15): 4785–90.

Cirelli Chiara, and Giulio Tononi. 2008. “Is Sleep Essential?” PLoS Biology. https://doi.org/10.1371/journal.pbio.0060216.

Cohen, Dror, Bruno van Swinderen, and Naotsugu Tsuchiya. 2018. “Isoflurane Impairs Low-Frequency Feedback but Leaves High-Frequency Feedforward Connectivity Intact in the Fly Brain.” eNeuro 5 (1). https://doi.org/10.1523/ENEURO.0329-17.2018.

Cohen, Dror, and Naotsugu Tsuchiya. 2018. “The Effect of Common Signals on Power, Coherence and Granger Causality: Theoretical Review, Simulations, and Empirical Analysis of Fruit Fly LFPs Data.” Frontiers in Systems Neuroscience 12 (July): 30.

Cohen, Dror, Oressia H. Zalucki, Bruno van Swinderen, and Naotsugu Tsuchiya. 2016. “Local Versus Global Effects of Isoflurane Anesthesia on Visual Processing in the Fly Brain.” eNeuro 3 (4). https://doi.org/10.1523/ENEURO.0116-16.2016.

Cortes, Corinna, and Vladimir Vapnik. 1995. “Support-Vector Networks.” Machine Learning 20 (3): 273–97.

Dag, Ugur, Zhengchang Lei, Jasmine Q. Le, Allan Wong, Daniel Bushey, and Krystyna Keleman. 2019. “Neuronal Reactivation during Post-Learning Sleep Consolidates Long-Term Memory in.” eLife 8 (February). https://doi.org/10.7554/eLife.42786.

Delorme, Arnaud, and Scott Makeig. 2004. “EEGLAB: An Open Source Toolbox for Analysis of Single-Trial EEG Dynamics Including Independent Component Analysis.” Journal of Neuroscience Methods 134 (1): 9– 21.

Dement, W., and N. Kleitman. 1957. “Cyclic Variations in EEG during Sleep and Their Relation to Eye Movements, Body Motility, and Dreaming.” Electroencephalography and Clinical Neurophysiology 9 (4): 673–90.

Donlea, Jeffrey M., Narendrakumar Ramanan, and Paul J. Shaw. 2009. “Use-Dependent Plasticity in Clock Neurons Regulates Sleep Need in Drosophila.” Science 324 (5923): 105–8.

Donlea, Jeffrey M., Matthew S. Thimgan, Yasuko Suzuki, Laura Gottschalk, and Paul J. Shaw. 2011. “Inducing Sleep by Remote Control Facilitates Memory Consolidation in Drosophila.” Science 332 (6037): 1571–76.

Eban-Rothschild, Ada D., and Guy Bloch. 2008. “Differences in the Sleep Architecture of Forager and Young honeybees(*Apis Mellifera*).” Journal of Experimental Biology. https://doi.org/10.1242/jeb.016915.

Faville, R., B. Kottler, G. J. Goodhill, P. J. Shaw, and B. van Swinderen. 2015. “How Deeply Does Your Mutant Sleep? Probing Arousal to Better Understand Sleep Defects in Drosophila.” Scientific Reports 5 (February): 8454.

Flood, Thomas F., Shinya Iguchi, Michael Gorczyca, Benjamin White, Kei Ito, and Motojiro Yoshihara. 2013. “A Single Pair of Interneurons Commands the Drosophila Feeding Motor Program.” Nature. https://doi.org/10.1038/nature12208.

Flores-Valle, Andres, and Johannes D. Seelig. 2022. “Dynamics of a Sleep Homeostat Observed in Glia during Behavior.” bioRxiv. https://doi.org/10.1101/2022.07.07.499175.

Fox, John, and Sanford Weisberg. 2018. An R Companion to Applied Regression. SAGE Publications. Fulda, Stephany, Christoph P. N. Romanowski, Andreas Becker, Thomas C. Wetter, Mayumi Kimura, and Thomas Fenzel. 2011. “Rapid Eye Movements during Sleep in Mice: High Trait-like Stability Qualifies Rapid Eye Movement Density for Characterization of Phenotypic Variation in Sleep Patterns of Rodents.” BMC Neuroscience 12 (November): 110.

Gilestro, Giorgio F., Giulio Tononi, and Chiara Cirelli. 2009. “Widespread Changes in Synaptic Markers as a Function of Sleep and Wakefulness in Drosophila.” Science 324 (5923): 109–12.

Grabowska, Martyna J., Rhiannon Jeans, James Steeves, and Bruno van Swinderen. 2020. “Oscillations in the Central Brain of Are Phase Locked to Attended Visual Features.” Proceedings of the National Academy of Sciences of the United States of America 117 (47): 29925–36.

Gramfort, Alexandre, Martin Luessi, Eric Larson, Denis A. Engemann, Daniel Strohmeier, Christian Brodbeck, Roman Goj, et al. 2013. “MEG and EEG Data Analysis with MNE-Python.” Frontiers in Neuroscience 7 (December): 267.

Hendricks, J. C., S. M. Finn, K. A. Panckeri, J. Chavkin, J. A. Williams, A. Sehgal, and A. I. Pack. 2000. “Rest in Drosophila Is a Sleep-like State.” Neuron 25 (1): 129–38.

Iglesias, Teresa L., Jean G. Boal, Marcos G. Frank, Jochen Zeil, and Roger T. Hanlon. 2019. “Cyclic Nature of the REM Sleep-like State in the Cuttlefish Sepia Officinalis.” The Journal of Experimental Biology 222 (1): jeb174862.

Jaggard, James B., Gordon X. Wang, and Philippe Mourrain. 2021. “Non-REM and REM/paradoxical Sleep Dynamics across Phylogeny.” Current Opinion in Neurobiology 71 (December): 44–51.

Jenett, Arnim, Gerald M. Rubin, Teri-T B. Ngo, David Shepherd, Christine Murphy, Heather Dionne, Barret D. Pfeiffer, et al. 2012. “A GAL4-Driver Line Resource for Drosophila Neurobiology.” Cell Reports 2 (4): 991–1001.

Kain, Pinky, and Anupama Dahanukar. 2015. “Secondary Taste Neurons That Convey Sweet Taste and Starvation in the Drosophila Brain.” Neuron 85 (4): 819–32.

Kaiser, Walter, and Jana Steiner-Kaiser. 1983. “Neuronal Correlates of Sleep, Wakefulness and Arousal in a Diurnal Insect.” Nature 301 (5902): 707–9.

Kim, Heesoo, Colleen Kirkhart, and Kristin Scott. 2017. “Long-Range Projection Neurons in the Taste Circuit of Drosophila.” eLife 6 (February). https://doi.org/10.7554/eLife.23386.

Kirszenblat, Leonie, and Bruno van Swinderen. 2015. “The Yin and Yang of Sleep and Attention.” Trends in Neurosciences 38 (12): 776–86.

Kuznetsova, Alexandra, Per B. Brockhoff, and Rune H. B. Christensen. 2017. “lmerTest Package: Tests in Linear Mixed Effects Models.” Journal of Statistical Software 82 (December): 1–26.

Leung, Angus, Dror Cohen, Bruno van Swinderen, and Naotsugu Tsuchiya. 2021. “Integrated Information Structure Collapses with Anesthetic Loss of Conscious Arousal in Drosophila Melanogaster.” PLoS Computational Biology 17 (2): e1008722.

Lin, Jian-Sheng, Pertti Panula, and Maria Beatrice Passani. 2015. Histamine in the Brain. Frontiers Media SA.

Mathis, Alexander, Pranav Mamidanna, Kevin M. Cury, Taiga Abe, Venkatesh N. Murthy, Mackenzie Weygandt Mathis, and Matthias Bethge. 2018. “DeepLabCut: Markerless Pose Estimation of User-Defined Body Parts with Deep Learning.” Nature Neuroscience 21 (9): 1281–89.

Miller, Patrick J. O., Kagari Aoki, Luke E. Rendell, and Masao Amano. 2008. “Stereotypical Resting Behavior of the Sperm Whale.” Current Biology: CB 18 (1): R21–23.

Mu, Laiyong, Kei Ito, Jonathan P. Bacon, and Nicholas J. Strausfeld. 2012. “Optic Glomeruli and Their Inputs in Drosophila Share an Organizational Ground Pattern with the Antennal Lobes.” The Journal of Neuroscience: The Official Journal of the Society for Neuroscience 32 (18): 6061–71.

Muñoz, Roberto N., Angus Leung, Aidan Zecevik, Felix A. Pollock, Dror Cohen, Bruno van Swinderen, Naotsugu Tsuchiya, and Kavan Modi. 2020. “General Anesthesia Reduces Complexity and Temporal Asymmetry of the Informational Structures Derived from Neural Recordings in *Drosophila*.” Physical Review Research 2 (2): 023219.*aaaa*

Nitz, Douglas A., Bruno van Swinderen, Giulio Tononi, and Ralph J. Greenspan. 2002. “Electrophysiological Correlates of Rest and Activity in Drosophila Melanogaster.” Current Biology: CB 12 (22): 1934–40.

Ostrovsky, Aaron, Sebastian Cachero, and Gregory Jefferis. 2013. “Clonal Analysis of Olfaction in Drosophila: Image Registration.” Cold Spring Harbor Protocols 2013 (4): 347–49.

Paulk, AngeliqDue C., Leonie Kirszenblat, Yanqiong Zhou, and Bruno van Swinderen. 2015. “Closed-Loop Behavioral Control Increases Coherence in the Fly Brain.” The Journal of Neuroscience: The Official Journal of the Society for Neuroscience 35 (28): 10304–15.

Paulk, Angelique C., Yanqiong Zhou, Peter Stratton, Li Liu, and Bruno van Swinderen. 2013. “Multichannel Brain Recordings in Behaving Drosophila Reveal Oscillatory Activity and Local Coherence in Response to Sensory Stimulation and Circuit Activation.” Journal of Neurophysiology 110 (7): 1703–21.

Raccuglia, Davide, Sheng Huang, Anatoli Ender, M-Marcel Heim, Desiree Laber, Raquel Suárez-Grimalt, Agustin Liotta, Stephan J. Sigrist, Jörg R. P. Geiger, and David Owald. 2019. “Network-Specific Synchronization of Electrical Slow-Wave Oscillations Regulates Sleep Drive in Drosophila.” Current Biology: CB 29 (21): 3611–21.e3.

Raccuglia, Davide, Raquel Suárez-Grimalt, Laura Krumm, Cedric B. Brodersen, Anatoli Ender, Sridhar R. Jagannathan, York Winter, et al. 2022. “Coherent Multi-Level Network Oscillations Create Neural Filters to Favor Quiescence over Navigation in Drosophila.” bioRxiv. https://doi.org/10.1101/2022.03.11.483976.

Rasch Björn, and Jan Born. 2013. “About Sleep’s Role in Memory.” Physiological Reviews. https://doi.org/10.1152/physrev.00032.2012.

Rößler, Daniela C., Kris Kim, Massimo De Agrò, Alex L. Jordan, Cosmas Giovanni Galizia, and Paul S. Shamble. 2022. Regularly Occurring Bouts of Retinal Movements Suggest an REM Sleep-like State in Jumping Spiders.

Sauer, S., M. Kinkelin, E. Herrmann, and W. Kaiser. 2003. “The Dynamics of Sleep-like Behaviour in Honey Bees.” *Journal of Comparative Physiology. A, Neuroethology, Sensory*, Neural, and Behavioral Physiology 189 (8): 599–607.

Scheffer, Louis K., C. Shan Xu, Michal Januszewski, Zhiyuan Lu, Shin-Ya Takemura, Kenneth J. Hayworth, Gary B. Huang, et al. 2020. “A Connectome and Analysis of the Adult Drosophila Central Brain,” September. https://doi.org/10.7554/eLife.57443.

Shafer, O. T., and A. C. Keene. 2021. “The Regulation of Drosophila Sleep.” Current Biology: CB 31 (1). https://doi.org/10.1016/j.cub.2020.10.082.

Shaw, Paul J., Giulio Tononi, Ralph J. Greenspan, and Donald F. Robinson. 2002. “Stress Response Genes Protect against Lethal Effects of Sleep Deprivation in Drosophila.” Nature 417 (6886): 287–91.

Shaw, P. J., C. Cirelli, R. J. Greenspan, and G. Tononi. 2000. “Correlates of Sleep and Waking in Drosophila Melanogaster.” Science 287 (5459): 1834–37.

Siegel, J. M. 2001. “The REM Sleep-Memory Consolidation Hypothesis.” Science 294 (5544): 1058–63.

Stanhope, Bethany A., James B. Jaggard, Melanie Gratton, Elizabeth B. Brown, and Alex C. Keene. 2020. “Sleep Regulates Glial Plasticity and Expression of the Engulfment Receptor Draper Following Neural Injury.” Current Biology*: CB* 30 (6): 1092–1101.e3.

Swinderen, Bruno van, and Ralph J. Greenspan. 2003. “Salience Modulates 20-30 Hz Brain Activity in Drosophila.” Nature Neuroscience 6 (6): 579–86.

van Swinderen, B., D. A. Nitz, and R. J. Greenspan. 2004. “Uncoupling of Brain Activity from Movement Defines Arousal States in Drosophila.” Current Biology: CB 14 (2): 81–87.

Tainton-Heap, Lucy A. L., Leonie C. Kirszenblat, Eleni T. Notaras, Martyna J. Grabowska, Rhiannon Jeans, Kai Feng, Paul J. Shaw, and Bruno van Swinderen. 2021. “A Paradoxical Kind of Sleep in Drosophila Melanogaster.” Current Biology: CB 31 (3): 578–90.e6.

Tononi, Giulio, and Chiara Cirelli. 2014. “Sleep and the Price of Plasticity: From Synaptic and Cellular Homeostasis to Memory Consolidation and Integration.” Neuron 81 (1): 12–34.

Troup, Michael, Lucy A. L. Tainton-Heap, and Bruno van Swinderen. 2023. “Neural Ensemble Fragmentation in the Anesthetized Brain.” The Journal of Neuroscience: The Official Journal of the Society for Neuroscience 43 (14): 2537–51.

Troup, Michael, Melvyn Hw Yap, Chelsie Rohrscheib, Martyna J. Grabowska, Deniz Ertekin, Roshini Randeniya, Benjamin Kottler, et al. 2018. “Acute Control of the Sleep Switch in Reveals a Role for Gap Junctions in Regulating Behavioral Responsiveness.” eLife 7 (August). https://doi.org/10.7554/eLife.37105.

Van De Poll, Matthew N., and Brunos van Swinderen. 2021. “Balancing Prediction and Surprise: A Role for Active Sleep at the Dawn of Consciousness?” Frontiers in Systems Neuroscience 15 (November): 768762.

Walker, Matthew P., and Robert Stickgold. 2004. “Sleep-Dependent Learning and Memory Consolidation.” Neuron 44 (1): 121–33.

Wiggin, Timothy D., Patricia R. Goodwin, Nathan C. Donelson, Chang Liu, Kien Trinh, Subhabrata Sanyal, and Leslie C. Griffith. 2020. “Covert Sleep-Related Biological Processes Are Revealed by Probabilistic Analysis in.” Proceedings of the National Academy of Sciences of the United States of America 117 (18): 10024–34.

Xie, Lulu, Hongyi Kang, Qiwu Xu, Michael J. Chen, Yonghong Liao, Meenakshisundaram Thiyagarajan, John O’Donnell, et al. 2013. “Sleep Drives Metabolite Clearance from the Adult Brain.” Science 342 (6156): 373–77.

Yap, Melvyn H. W., Martyna J. Grabowska, Chelsie Rohrscheib, Rhiannon Jeans, Michael Troup, Angelique C. Paulk, Bart van Alphen, Paul J. Shaw, and Bruno van Swinderen. 2017. “Oscillatory Brain Activity in Spontaneous and Induced Sleep Stages in Flies.” Nature Communications 8 (1): 1815.

